# Multiple *pals* gene modules control a balance between immunity and development in *Caenorhabditis elegans*

**DOI:** 10.1101/2023.01.15.524171

**Authors:** Vladimir Lažetić, Michael J. Blanchard, Theresa Bui, Emily R. Troemel

**Affiliations:** School of Biological Sciences, University of California, San Diego, La Jolla, California, United States

## Abstract

The immune system continually battles against pathogen-induced pressures, which often leads to the evolutionary expansion of immune gene families in a species-specific manner. For example, the *pals* gene family expanded to 39 members in the *Caenorhabditis elegans* genome, in comparison to a single mammalian *pals* ortholog. Our previous studies have revealed that two members of this family, *pals-22* and *pals-25*, act as antagonistic paralogs to control the Intracellular Pathogen Response (IPR). The IPR is a protective transcriptional response, which is activated upon infection by two molecularly distinct natural intracellular pathogens of *C. elegans* – the Orsay virus and the fungus *Nematocida parisii* from the microsporidia phylum. In this study, we identify a previously uncharacterized member of the *pals* family, *pals-17*, as a newly described negative regulator of the IPR. *pals-17* mutants show constitutive upregulation of IPR gene expression, increased immunity against intracellular pathogens, as well as impaired development and reproduction. We also find that two other previously uncharacterized *pals* genes, *pals-20* and *pals-16*, are positive regulators of the IPR, acting downstream of *pals-17*. These positive regulators reverse the effects caused by the loss of *pals-17* on IPR gene expression, immunity and development. We show that the negative IPR regulator protein PALS-17 and the positive IPR regulator protein PALS-20 colocalize inside intestinal epithelial cells, which are the sites of infection for IPR-inducing pathogens. In summary, our study demonstrates that several *pals* genes from the expanded *pals* gene family act as ON/OFF switch modules to regulate a balance between organismal development and immunity against natural intracellular pathogens in *C. elegans*.

**AUTHOR SUMMARY:** Immune responses to pathogens induce extensive rewiring of host physiology. In the short term, these changes are generally beneficial as they can promote resistance against infection. However, prolonged activation of immune responses can have serious negative consequences on host health, including impaired organismal development and fitness. Therefore, the balance between activating the immune system and promoting development must be precisely regulated. In this study, we used genetics to identify a gene in the roundworm *Caenorhabditis elegans* called *pals-17* that acts as a repressor of the Intracellular Pathogen Response (IPR), a defense response against viral and microsporidian infections. We also found that *pals-17* is required for the normal development of these animals. Furthermore, we identified two other *pals* genes, *pals-20* and *pals-16*, as suppressors of *pals-17* mutant phenotypes. Finally, we found that PALS-17 and PALS-20 proteins colocalize inside intestinal cells, where viruses and microsporidia invade and replicate in the host. Taken together, our study demonstrates a balance between organismal development and immunity that is regulated by several genetic ON/OFF switch ‘modules’ in *C. elegans*.

## INTRODUCTION

Organismal survival and evolutionary success require organisms to balance the demands of defense against infection, with the demands of development and reproduction. Research from several organisms has demonstrated that there can be trade-offs between the demands for immunity and the demands for growth. For example, enhancing immune responses in plants often leads to impaired growth and lowered crop yields, a trade-off that is regulated by secondary messengers and phytohormones (1-3). Compared to plants, less is known about the mechanisms by which animals tune the balance between growth and defense in order to promote their overall survival and reproduction.

Recently, we described a genetic switch in the nematode *C. elegans* that regulates a balance between growth and immunity (4, 5). Through forward genetic screening, we found that the *pals-22* gene represses a set of immunity-related genes called the Intracellular Pathogen Response (IPR) genes, which are activated upon infection by natural intracellular pathogens of the *C. elegans* intestine – the Orsay virus and the fungus *Nematocida parisii* from the microsporidia phylum (6-10). In the absence of infection, mutations in *pals-22* lead to constitutive expression of IPR genes, increased immunity against intracellular pathogens, and slowed growth. Through suppressor screens, we found that mutations in the *pals-25* gene reverse *pals-22* mutant phenotypes back to wild-type, indicating that *pals-25* is required for mediating the effects caused by the loss of *pals-22* (4). Thus, the antagonistic paralogs *pals-25* and *pals-22* comprise an ON/OFF switch module that increases immunity with an associated cost of slowed growth.

Genes in the *pals* family encode proteins that contain the ALS2CR12 domain, a divergent amino acid signature of unknown biochemical function (6, 11). Genes associated with immunity often undergo species-specific expansion (12, 13), and accordingly, there are 39 *pals* genes found in *C. elegans*, and only a single *pals* gene found each in mice and humans, with no reported function in those organisms. The *C. elegans pals-22* and *pals-25* genes are co-expressed in an operon, and most *pals* genes in *C. elegans* are clustered together in the genome, suggesting that they resulted from gene duplication events. The *pals* gene family exhibits sequence divergence and species-specific changes in gene number within species of the *Caenorhabditis* genus. While *C. elegans* has 39 *pals* genes, there are 18 *pals* genes in *C. remanei*, and only 8 *pals* genes each in *C. briggsae* and *C. brenneri (11)*. Of the 39 *pals* genes in *C. elegans*, 26 of them are upregulated by IPR triggers, including the *pals-5* gene, which is commonly used as a reporter for the IPR (*pals-5*p::GFP) (6). Notably, *pals-22* and *pals-25* are not induced by infection, leading to the hypothesis that non-induced *pals* genes serve as regulators of induced *pals* genes.

Here, through a forward genetic screen for mutants that have constitutive *pals-5*p::GFP expression in the absence of infection, we identify *pals-17* as a new negative regulator of IPR gene expression, acting independently of *pals-22*. Through targeted RNA interference (RNAi) screening of non-induced *pals* genes, we identify *pals-20* and *pals-16* as two positive regulators that act downstream of *pals-17* to induce IPR gene expression. Notably, *pals-16, pals-17* and *pals-20* are not induced by infection, similar to *pals-22* and *pals-25*, supporting the hypothesis that non-induced *pals* genes are regulators of induced *pals* genes. We show that PALS-17 and PALS-20 proteins are co-expressed in the intestine. We also find that point mutations in *pals-17* lead to slowed growth and increased immunity against natural intracellular pathogens of the intestine, while a full *pals-17* deletion leads to developmental arrest. The immunity and growth phenotypes of *pals-17* mutants are dependent on the positive IPR regulators *pals-16* and *pals-20*. Consistent with *pals-20* being an activator of the IPR, we find that overexpression of *pals-20* in a wild-type background promotes resistance to viral infection. Altogether, these results identify PALS-17 as a negative regulator of the IPR similar to PALS-22, and PALS-16 and PALS-20 as positive regulators of the IPR similar to PALS-25. These findings indicate there are two separate modules of IPR regulators including the PALS-22/25 module and PALS-17/16/20 module, highlighting how PALS proteins in *C. elegans* have extensive control over the balance between growth and immunity, which is critical for organismal success.

## RESULTS

### *pals-17* is a negative regulator of the IPR and an essential gene

To identify novel regulators of the IPR, we performed a forward genetic screen for mutations that cause constitutive expression of the transcriptional IPR reporter *pals-5*p::GFP (*jyIs8*). Here, we mutagenized a strain that has loss-of-function mutations in *pals-22* and *pals-25*, two previously identified regulators of the IPR. We performed the screen in this double mutant to prevent isolation of new alleles of *pals-22* or *pals-25*. Here we identified the allele *jy74*, which exhibits constitutive expression of the *pals-5*p::GFP reporter in intestinal epithelial cells (Fig 1A-D, S1 Table). Whole-genome sequencing of *jy74* mutants identified a mutation in the gene *pals-17* (Fig 1E). Specifically, the *jy74* allele, with guanine 325 mutated into adenine, is predicted to affect splicing between the third and fourth exons of *pals-17*. *pals-17*, like *pals-22* and *pals-25*, belongs to the clade of *pals* genes that are not induced upon intracellular infection (4-7). To confirm that the mutation in *pals-17* is responsible for the upregulated expression of the *pals-5*p::GFP reporter, we introduced a fosmid containing a wild-type copy of *pals-17* into *pals-22(-) pals-25(-) pals-17(jy74)* mutants. Here, we observed a decrease in *pals-5*p::GFP expression in animals carrying the *pals-17* fosmid array compared to their siblings without the array. This observation indicates that loss of *pals-17* function is responsible for constitutive *pals-5*p::GFP expression in *jy74* mutants (Fig 1F).

**Fig 1.**
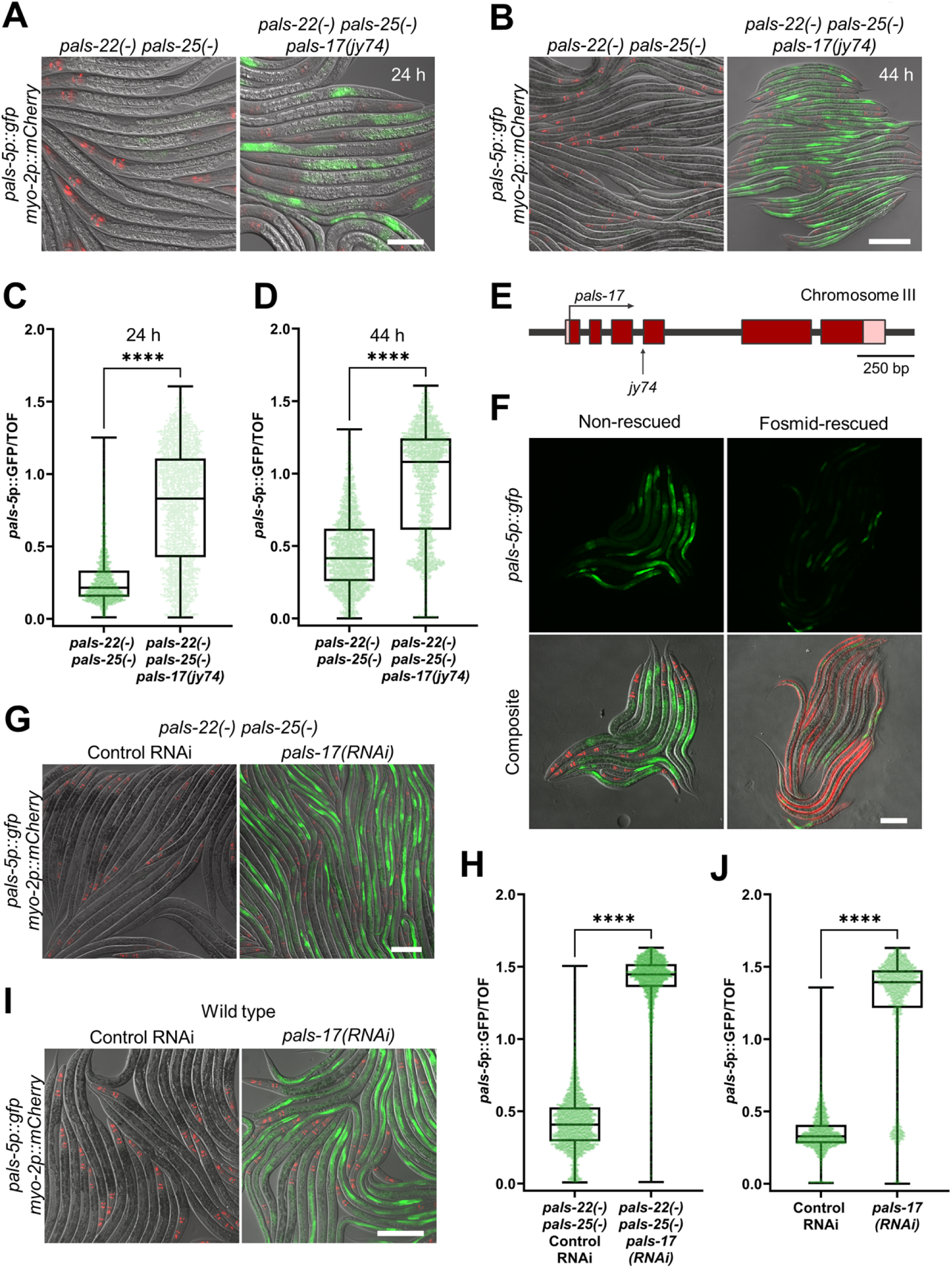
*pals-17* is a negative regulator of *pals-5*p::GFP expression. (A-D) *pals-17(jy74)* mutants show increased *pals-5*p::GFP expression. Representative images of synchronized populations of the control *pals-22(-) pals-25(-)* strain and *pals-22(-) pals-25(-) pals-17(jy74)* mutants grown for 24 h (A) and 44 h (B) at 20 °C from L1 stage. Red, green and DIC channels were merged. *myo-2p::mCherry* co-injection marker is shown in red. Scale bars, 50 µm (A) and 200 µm (B). Levels of *pals-5*p::GFP expression standardized to time-of-flight (TOF) are indicated by box-and-whisker plots for the control strain and *pals-17(jy74)* mutants grown for 24 h (C) and 44 h (D) at 20 °C from L1 stage. (E) *pals-17* gene structure. Exons are indicated with dark red boxes, 5’ and 3’ UTRs are shown with light red boxes. *jy74* mutation indicated with vertical arrow, alters the predicted splice site for the fourth exon. The horizontal arrow indicates the direction of transcription. (F) A rescuing fosmid containing wild-type *pals-17,* injected together with the *myo-3p::mCherry* body-wall muscle marker, reduces *pals-5*p::GFP expression in *pals-17(jy74)* mutant animals (right), compared to their *pals-17(jy74)* mutant siblings that do not contain the fosmid (left). Red, green and DIC channels were merged in the composite images. *myo-2*p::mCherry and *myo-3*p::mCherry co-injection markers are shown in red. Scale bar, 100 µm. (G-J) *pals-17* RNAi increases *pals-5*p::GFP expression in *pals-22(-) pals-25(-)* mutants and wild-type animals. Representative images of synchronized L4 stage populations of the *pals-22(-) pals-25(-)* (G) and wild-type animals (I) grown on the control RNAi vector and *pals-17* RNAi plates. Red, green and DIC channels were merged. *myo-2*p::mCherry co-injection marker is shown in red. Scale bars, 50 µm. (H, J) Box-and-whisker plots of *pals-5*p::GFP expression levels standardized to TOF following control RNAi and *pals-17* RNAi treatments of *pals-22(-) pals-25(-)* mutants (H) and wild-type animals (J). (C, D, H, J) Lines in box-and-whisker plots represent the medians, box bounds indicate 25^th^ and 75^th^ percentiles, and whiskers extend from the box bounds to the minimum and maximum values. Green dots represent individual values for each animal; 600 (C), 800 (D), 1200 (H) and 750 animals (J) were analyzed per strain per each of the three replicates. A Kolmogorov-Smirnov test was used to calculate p-values; **** p < 0.0001.

To further confirm the role of *pals-17* in *pals-5*p::GFP regulation, we treated *pals-22 pals-25* double mutant animals with *pals-17* RNAi. Upon knockdown of *pals-17*, we observed increased expression of the *pals-5*p::GFP reporter, consistent with *pals-17* acting as a negative regulator (Fig 1G and 1H, S1 Table). Next, we investigated whether loss of *pals-17* induces *pals-5*p::GFP expression in the presence of functional *pals-22* and *pals-25*. Here, we performed the same RNAi analysis in a wild-type background and observed a similar phenotype; *pals-5*p::GFP expression was significantly induced following *pals-17* RNAi treatment (Fig 1I and 1J, S1 Table). Thus, loss of *pals-17* causes upregulation of *pals-5*p::GFP expression in both a wild-type background and a *pals-22 pals-25* mutant background, suggesting that *pals-17* acts independently of *pals-22* and *pals-25*.

In addition to *pals-5*p::GFP expression, we also examined if the loss of *pals-17* upregulates ZIP-1::GFP expression. In recent work, we demonstrated that ZIP-1 is a transcription factor that promotes immunity against intracellular pathogens (14). All previously described triggers require ZIP-1 for induction of the *pals-5*p::GFP reporter. Here, we found that *pals-17* RNAi induces expression of ZIP-1::GFP in intestinal nuclei (Fig S1), which also occurs in response to several other IPR triggers (5, 14).

To determine the null phenotype of *pals-17*, the entire coding region of *pals-17* was deleted to generate the *pals-17(Δ)* allele (S2A Fig). We found that these *pals-17(Δ)* null mutants failed to develop past an early larval stage (approximately first larval stage L1 to second larval stage L2), indicating that *pals-17* is an essential gene (S2B Fig). To quantify this effect, we measured the body length of progeny from a balanced *pals-17(Δ)/sC1(s2023)[dpy-1(s2170) umnIs21]* strain and observed significantly smaller body size for *pals-17* homozygotes in comparison to their *pals-17* heterozygous siblings, to their Dpy (wildtype *pals-17*) siblings, and to wild-type control animals (S2C Fig, S1 Table). Notably, we observed that this larval arrest phenotype is 100% penetrant, as all *pals-17(Δ)* homozygotes failed to develop into adults (60/60 from three replicates). In summary, our analyses demonstrate that *pals-17* is a negative regulator of the IPR reporter, and is essential for *C. elegans* development.

### Loss of *pals-20* partially suppresses *pals-17* mutant phenotypes

The analysis above indicates that wild-type *pals-17* acts as a repressor of IPR gene expression, similar to wild-type *pals-22*. Activation of the IPR in *pals-22* mutants requires the *pals-25* gene, which is just downstream of *pals-22* in the genome as part of an operon (4). Therefore, we hypothesized that a gene that is positioned immediately downstream of *pals-17* might antagonize *pals-17* function, similar to how *pals-25* antagonizes *pals-22* function. Because *pals-20* is the closest neighbor of *pals-17* in the genome (although they are not in an operon), we performed RNAi against *pals-20* in *pals-22(-) pals-25(-) pals-17(jy74)* mutants and examined *pals-5*p::GFP expression. Notably, we observed a slight decrease in GFP reporter expression in *pals-20(RNAi)* animals compared to control RNAi, indicating that *pals-20* may be an activator of the IPR (Fig 2A).

**Fig 2.**
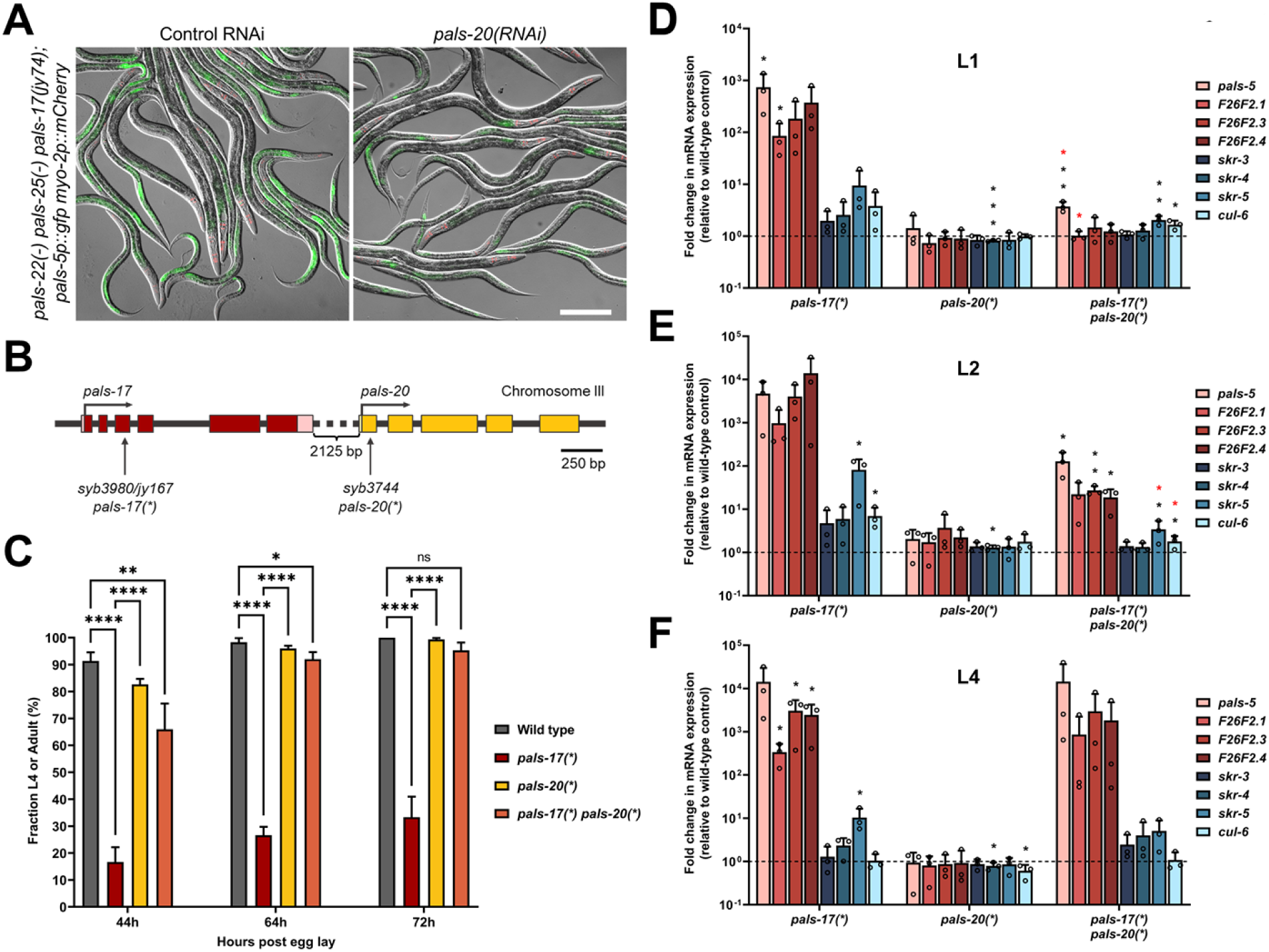
*pals-17* mutant phenotypes are partially dependent on *pals-20*. (A) Representative images of *pals-5*p::GFP expression in *pals-22(-) pals-25(-) pals-17(jy74)* mixed stage animals on control and *pals-20* RNAi plates. Green, red and DIC channels were merged. *myo-2*p::mCherry co-injection marker is shown in red. Scale bar, 200 µm. (B) *pals-17* and *pals-20* gene structures. *pals-17* exons are indicated with dark red boxes, 5’ and 3’ UTRs are shown with light red boxes. *pals-20* exons are indicated with yellow boxes. Vertical arrows indicate the positions of the point mutations that create premature stop codons in *pals-17* and *pals-20*. Horizontal arrows indicate the directions of transcription. (C) Fractions of animals reaching the L4 or adult stage at 44, 64 and 72 hours after eggs were laid. Results are averages of three independent experimental replicates, with 100 animals in each replicate at each time point. Error bars are standard deviations (SD). Ordinary one-way ANOVA test was used to calculate p-values; **** p < 0.0001; ** p < 0.01; * p < 0.05; ns indicates no significant difference. (D-F) *pals-20* mutation suppresses increased expression of IPR genes in the *pals-17* mutant background at early life stages. qRT-PCR of IPR gene expression shown as the mean fold changes relative to the wild-type control at L1 (D), L2 (E) and L4 stage (F). Error bars are SD. A one-tailed t-test was used to calculate p-values; black asterisks represent a significant difference between the labeled sample and the wild-type control; red asterisks represent a significant difference between *pals-17(*)* and *pals-17(*) pals-20(*)* samples; *** p < 0.001; ** p < 0.01; * p < 0.05; p-values higher than 0.05 are not labeled.

To further investigate the function of *pals-20*, a mutant allele that contains a premature stop codon in the first exon of *pals-20* was generated (*pals-20(*)*); see Materials and Methods for more information) (Fig 2B). Similar to *pals-25* loss-of-function alleles, the *pals-20(*)* loss-of-function allele had no apparent phenotype in a wild-type background. Next, we investigated whether *pals-20(*)* would suppress *pals-17* mutant phenotypes. Because a full deletion of *pals-17* led to larval arrest, we sought another *pals-17* allele, with the goal of creating a mutation that would cause a strong loss-of-function, but allow animals to develop into adulthood. To achieve this goal, the *pals-17(*)* allele was generated, which contains an early stop in the third exon in *pals-17*. This mutation causes slowed and asynchronous development, but allows animals to reach adulthood and reproduce (Fig 2B and 2C, S1 Table). Here we found that the *pals-20(*)* mutation substantially suppressed the developmental delay and asynchronous development of *pals-17* mutants. These *pals-17 pals-20* double mutants had slightly slower developmental rates than *pals-20* single mutants and the wild-type control, but they developed significantly faster than *pals-17* mutants (Fig 2C, S1 Table). To better understand the developmental delay of *pals-17* mutants and to determine conditions to better synchronize these animals for further analyses, we measured the gonad development of *pals-17* and *pals-20* mutant strains. Here, we found that gonad development in *pals-17* mutants is delayed and highly asynchronous, while *pals-17 pals-20* double mutants at the L2 stage have similar gonad areas to wild-type and *pals-20* single mutants (S3 Fig). Taken together, these results show that the loss of *pals-20* can suppress the developmental delay and asynchrony of *pals-17* mutants.

We next examined the endogenous mRNA expression of IPR genes at different developmental stages in *pals-17*, *pals-20*, and *pals-17 pals-20* double mutants. First, we performed qRT-PCR analysis in L1 animals. We found significant upregulation of IPR genes in the *pals-17* mutant background, which was almost completely suppressed in *pals-17 pals-20* double mutants (Fig 2D, S1 Table). *pals-20* single mutants had wild-type levels of IPR gene expression. Next, we found that the upregulated IPR gene expression observed in *pals-17* mutants was only partially suppressed in *pals-17 pals-20* double mutants at the L2 stage (Fig 2E, S1 Table). Finally, at the L4 stage, we surprisingly observed no suppression of IPR gene induction in *pals-17* mutants by loss of *pals-20* (Fig 2F, S1 Table). In summary, we found that endogenous mRNA levels of multiple IPR genes are highly induced in *pals-17* mutants from the L1 to the L4 stage. This induction is *pals-20*-dependent early in life, but less *pals-20*-dependent as animals develop into later larval stages.

### Loss of *pals-16* fully suppresses *pals-17* mutant phenotypes

Because IPR induction in *pals-17* mutants is dependent on the *pals-20* gene only in early life, we screened for a gene that plays a similar role to *pals-20* later in life. Here, we used RNAi to test several previously uncharacterized *pals* gene family members, focusing on those that are not strongly induced by intracellular infection (6, 11). These genes included *pals-1*, *pals-13*, *pals-16*, *pals-18*, *pals-19*, *pals-23*, *pals-24*, *pals-31*, *pals-34*, *pals-39* and *pals-40* (S4A Fig). In this assay, we used *pals-17 pals-20* double mutant animals that express *pals-5*p::GFP in the intestine at the adult stage and looked for an RNAi clone that would decrease GFP expression. Here we found that *pals-16* RNAi significantly decreased *pals-5*p::GFP expression (Fig 3A and 3B, S1 Table), suggesting that *pals-16* acts downstream of *pals-17* later in life.

**Fig 3.**
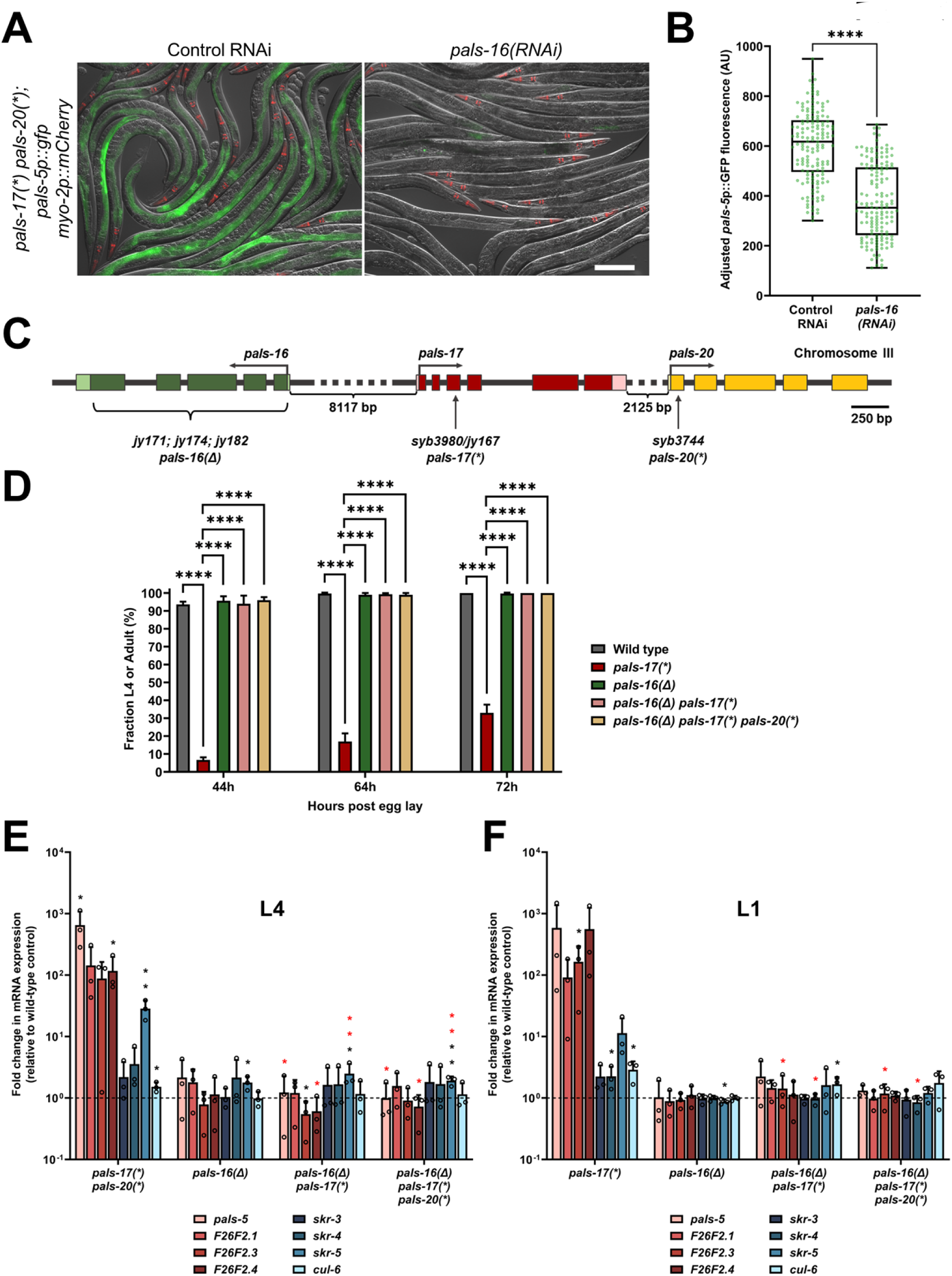
*pals-17* mutant phenotypes are fully *pals-16*-dependent. (A, B) *pals-16* RNAi decreases *pals-5*p::GFP expression in *pals-17 pals-20* mutants. (A) Representative images of synchronized *pals-17(*) pals-20(*)* mutants grown on control or *pals-16* RNAi plates for 72 h at 20 °C. Red, green and DIC channels were merged. *myo-2p::mCherry* co-injection marker is shown in red. Scale bar, 200 µm. (B) Box-and-whisker plot of *pals-5*p::GFP expression levels of (A) adjusted to background fluorescence for each sample. Line represents median, box bounds indicate 25^th^ and 75^th^ percentiles, and whiskers indicate minimum and maximum values. Green dots represent individual values for each animal; 50 animals were analyzed per strain per each of the three replicates. A Kolmogorov-Smirnov test was used to calculate p-values; **** p < 0.0001. AU = arbitrary units. (C) *pals-16*, *pals-17* and *pals-20* gene structures. *pals-16* exons indicated with dark green boxes, 5’ and 3’ UTRs with light green boxes. *pals-17* exons indicated with dark red boxes, 5’ and 3’ UTRs as light red boxes. *pals-20* exons indicated with yellow boxes. Bracket under green boxes demarks approximate area deleted in *jy171*, *jy174* and *jy182 pals-16* alleles (*pals-16(Δ)*). Vertical arrows indicate positions of point mutations that create premature stop codons in *pals-17* and *pals-20*. Horizontal arrows indicate directions of transcription. (D) Fractions of animals reaching L4 or adult stage at 44, 64 and 72 hours after eggs were laid. Results shown are the average of three independent experimental replicates, with 100 animals assayed in each replicate at each time point. Error bars are SD. An ordinary one-way ANOVA test was used to calculate p-values; **** p < 0.0001. (E, F) *pals-16* mutation suppresses increased expression of several IPR genes in the *pals-17(*)* and *pals-17(*) pals-20(*)* backgrounds. qRT-PCR measurements of gene expression are shown as the fold change relative to the wild-type control at L4 (E) and L1 stage (F). Mean values shown, error bars are SD. A one-tailed t-test was used to calculate p-values; black asterisks represent a significant difference between the labeled sample and wild-type control; red asterisks represent significant differences between *pals-17(*) pals-20(*)* and either *pals-16(Δ) pals-17(*)* or *pals-16(Δ) pals-17(*) pals-20(*)* (E), and significant differences between *pals-17(*)* and either *pals-16(Δ) pals-17(*)* or *pals-16(Δ) pals-17(*) pals-20(*)* (F); ** p < 0.01; * p < 0.05; p-values higher than 0.05 are not labeled.

To further examine this role of *pals-16*, we deleted the *pals-16* gene in wild-type animals, in *pals-17* mutants, and in *pals-17 pals-20* double mutants (Fig 3C). Here we found that the deletion of *pals-16*, called *pals-16(Δ),* was able to rescue the developmental defects of *pals-17* mutants. In contrast, *pals-16(Δ)* did not have any obvious effects in the other two backgrounds that have wild-type rates of development (Fig 3D, S1 Table). We then measured endogenous mRNA expression of IPR genes with qRT-PCR at L1 and L4 stages to test if and when *pals-16(Δ)* can suppress *pals-17*-dependent IPR induction. Here we found that the loss of *pals-16* leads to a decrease in mRNA levels of multiple IPR genes that are upregulated in *pals-17 pals-20* double mutants at the L4 stage (Fig 3E, S1 Table). Notably, we found that *pals-16(Δ)* is also able to suppress IPR gene induction in the *pals-17* single mutant background, independent of *pals-20*. Furthermore, *pals-16(Δ)* was able to suppress IPR gene expression in *pals-17* mutant animals at the L1 stage (Fig 3F, S1 Table). Taken together, our data demonstrate that *pals-16(Δ)* suppresses IPR induction in *pals-17* mutants throughout development and in a manner independent of *pals-20*.

Interestingly, PALS-20 and PALS-16 proteins cluster together with PALS-25 in a phylogenetic tree, due to similarities in amino acid sequence (S4A and S4B Fig). Specifically, PALS-20 and PALS-16 are 52% identical in amino acid sequence (S2 Table). Their similarities are also reflected in predicted AlphaFold protein structures (S4C Fig) (15). Computational models for PALS-20 and PALS-16 indicate that both proteins have four α-helix bundles in both their N- and C-termini, with a single long α-helix that connects them. Similarities of both PALS-20 and PALS-16 compared to PALS-17 are less pronounced, both in amino acid sequence (14-16% identical) and predicted structure (S4C Fig, S2 Table). While the predicted C-terminus of PALS-17 also has a four α-helix bundle, the predicted N-terminus of PALS-17 contains only two α-helices, and a longer intrinsically disordered region that connects the N-terminal α-helices to the central α-helix. Next, we compared these structural models with predicted structures for two previously described IPR regulators, PALS-25 and PALS-22 (S4D Fig). We found similarities among predicted structures of positive IPR regulators PALS-25, PALS-20, and PALS-16. All three structures have four α-helix bundles at both the N- and C-termini, and a long central α-helix. There were also some similarities in protein organization between the predictions for the two negative regulators of the IPR, PALS-17 and PALS-22. They both had an N-terminal two α-helix bundle, a long intrinsically disordered region, a central α-helix, and a C-terminal four α-helix bundle. In summary, this computational analysis suggests that there are similarities in the predicted PALS protein structures and their functions as either positive or negative regulators of the IPR.

### RNA sequencing analysis reveals a genome-wide picture of genes dependent on the *pals-16 pals-17 pals-20* regulatory module

Next, we used RNA sequencing (RNA-seq) to examine genome-wide transcriptional changes in *pals-17*, *pals-20*, and *pals-16* mutants. We performed our RNA-seq analysis at the L4 stage because this larval stage had the best RNA quality. First, we compared *pals-17* mutants to wild-type animals, growing them for different lengths of time in order to harvest both strains at approximately the L4 stage. Unfortunately, *pals-17* mutants have a high degree of individual variability in developmental time (S3 Fig); thus, it was not possible to obtain a pure population of L4 animals. Instead, we harvested a population of animals with the average stage of L4 that also contained some older and younger animals. In differential expression analysis of our RNA-seq data, we found that 3,001 genes were upregulated in *pals-17* mutants compared to wild-type animals, which significantly overlap with previously defined IPR genes (70/80 IPR genes) and genes upregulated in *pals-22(jy3)* mutants (262/457) (5) (Fig 4A, S1, S3 and S4 Tables). However, many of the differentially expressed genes in *pals-17* mutants could be due to the wide range of developmental ages found in a population of these mutants (S3 Fig). To address this issue, we analyzed mRNA expression of *pals-17 pals-20* mutants. These animals have upregulated IPR gene expression at the L4 stage similar to *pals-17* mutants, but they grow more synchronously within a population, at a rate comparable to wild-type animals (Fig 2, S3 Fig). We found that 190 genes were upregulated in *pals-17 pals-20* double mutants when compared to the wild type at L4. The majority of these genes were also upregulated in *pals-22* mutants (188/190) and they significantly overlapped with IPR genes (66/80 IPR genes) (Fig 4B, S1, S3 and S4 Tables). Furthermore, most of the genes upregulated in *pals-17 pals-20* double mutants (172/190) were also upregulated in *pals-17* single mutants (Fig 4C, S1 Table). To identify genes regulated by *pals-16*, we also performed RNA-seq on *pals-16 pals-17* double and *pals-16 pals-17 pals-20* triple mutant animals. Here we found that the upregulation of genes in *pals-17 pals-20* mutants was almost completely suppressed by the loss of *pals-16* (182/190) (Fig 4C, S1, S3 and S4 Tables).

**Fig 4.**
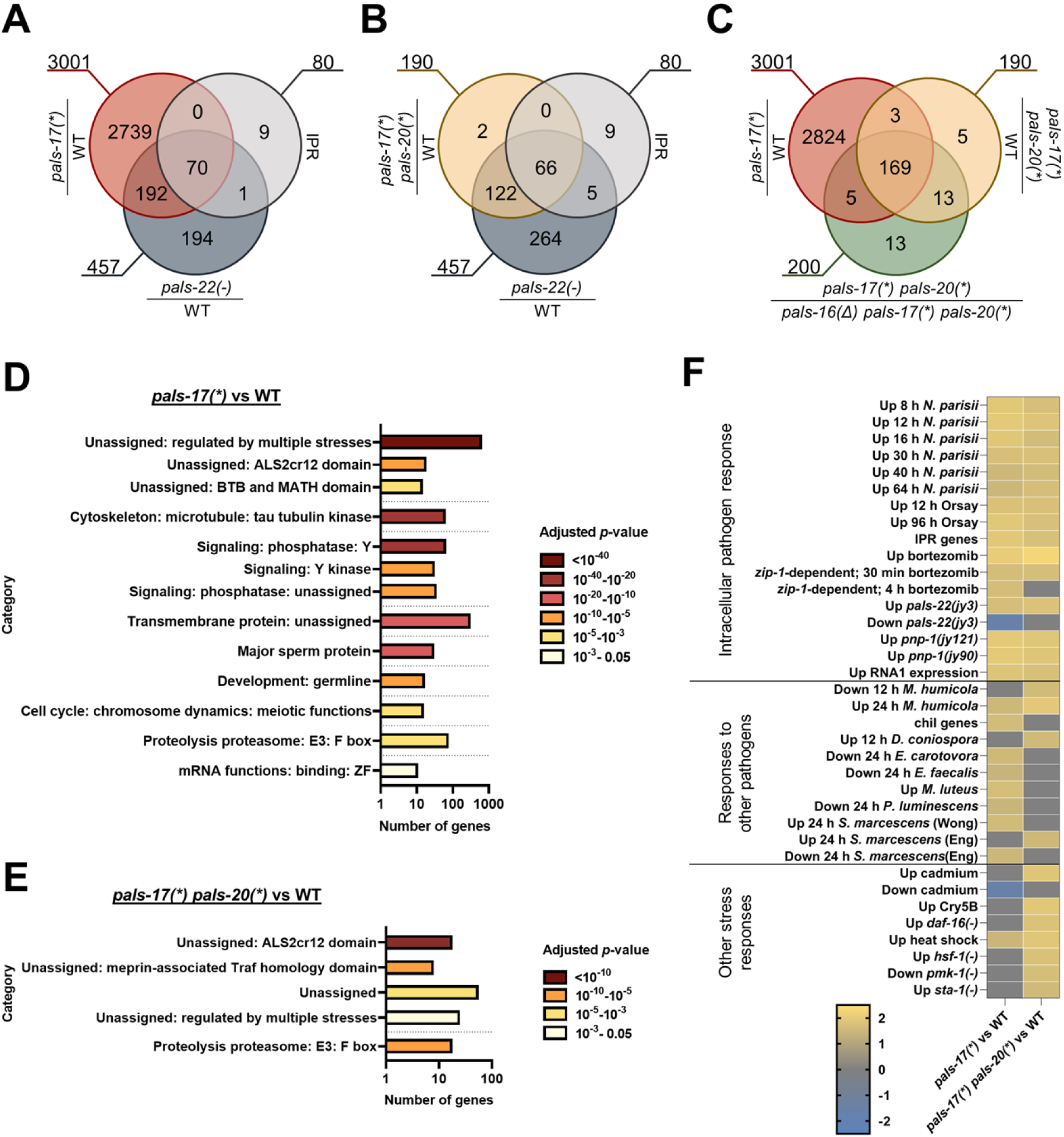
RNA-seq analyses demonstrate that *pals-17*, *pals-20* and *pals-16* regulate the expression of many genes at the L4 stage. (A-C) Venn diagram comparisons of genes upregulated in different genetic backgrounds. All RNAseq analyses were done in this study, except *pals-22(-)*-upregulated genes and IPR genes, which were determined in previous studies (4, 5). (A) *pals-17* is a negative regulator of IPR genes. Upregulated genes in *pals-17(*)* and *pals-22(-)* mutants as compared to wild-type (WT) animals, and to IPR genes. Upregulated genes in *pals-17(*)* mutants have significant overlap with those upregulated in *pals-22(-)* mutants (representation factor (rf) = 3.4; p < 5.788e-87) and with IPR genes (rf = 5.1; p < 2.155e-43). (B) IPR genes are still upregulated in *pals-17 pals-20* double mutant L4 animals. Upregulated genes *in pals-17(*) pals-20(*)* mutants as compared to WT have significant overlap with those upregulated in *pals-22(-)* mutants as compared to WT (rf = 38.1; p < 8.209e-314) and to IPR genes (rf = 76.5; p < 5.950e-121). (C) Upregulated genes in *pals-17(*)* mutants have significant overlap with those upregulated in *pals-17(*) pals-20(*)* mutants (rf = 5.3; p < 2.983e-111), both as compared to WT. Mutations in *pals-16* suppress genes upregulated in *pals-17* mutant backgrounds. Upregulated genes in *pals-17(*)* mutants when compared to WT show significant overlap with genes upregulated in *pals-17(*) pals-20(*)* mutants that are dependent on *pals-16* (rf = 5.1; p < 8.191e-106). Upregulated genes in *pals-17(*) pals-20(*)* mutants when compared to WT and *pals-16(Δ) pals-17(*) pals-20(*)* show significant overlap (rf = 84.3; p < 0.000e+00). (A-C) rf is the ratio of actual overlap to expected overlap where rf > 1 indicates overrepresentation and rf < 1 indicates underrepresentation. The numbers outside the circles represent the total number of genes in each dataset. (D, E) Enriched gene categories for *pals-17(*)*-dependent (D) and *pals-17(*) pals-20(*)*-dependent genes (E). Each category represents a biological process or a structure. The count of genes found in each category is indicated on the x-axis. Statistical significance for each category is indicated by the bar color; p-values are indicated in the legend on the right. p-values were determined using Bonferroni correction from the minimum hypergeometric scores calculated by the WormCat software. Source data (including p-values) are provided in S1 Table. Y = tyrosine; ZF = zinc finger. (F) Correlation of differentially expressed genes in *pals-17(*)* and *pals-17(*) pals-20(*)* mutants (in comparison to WT) with the genes that are differentially expressed during infection and exposure to other stressors. The correlation of gene sets was quantified as Normalized Enrichment Score (NES) as defined by GSEAPreranked module analysis. Yellow indicates a significant correlation of upregulated genes in *pals-17(*)* and/or *pals-17(*) pals-20(*)* mutants with the gene sets tested; grey indicates no significant correlation (p > 0.05 or False Discovery Rate > 0.25); blue indicates a significant correlation of downregulated genes in *pals-17(*)* and/or *pals-17(*) pals-20(*)* mutants with the gene sets tested. GSEA analysis of 93 gene sets tested and lists of genes are provided in S1 and S5 Tables.

To obtain insight into biological processes and cellular structures relevant to the genes upregulated in *pals-17* and *pals-17 pals-20* mutants, we performed analysis using the WormCat program, which is specifically designed to analyze *C. elegans* genomics data (16). We found that some of the genes upregulated in *pals-17* mutants belong to the category of genes that encode proteins with an ALS2CR12 domain – the defining hallmark of the PALS protein family (11), many of which are induced as part of the IPR (S4A Fig). Other categories included genes involved in protein degradation (F-box proteins), transcriptional regulation, reproduction, and signaling (Fig 4D, S1 Table). We confirmed the role of *pals-17* in reproduction in a brood size assay in which we found that *pals-17* mutants lay significantly fewer eggs. Furthermore, this phenotype was suppressed by the loss of *pals-20* (S5 Fig, S1 Table). The *pals-17 pals-20* RNA-seq dataset was enriched for ALS2CR12 domain and F-box categories, as well as multiple stress responses in the WormCat analysis (Fig 4E, S1 Table).

We next examined the correlation between *pals-17*- and *pals-17 pals-20*-dependent genes and gene sets that were previously associated with IPR activation, infection, and stress responses. We performed Gene Set Enrichment Analysis (GSEA) (17, 18) and found that *pals-17* and *pals-17 pals-20* mutants upregulate genes in common with infections by intracellular pathogens as well as known activators and regulators of the IPR (Fig 4F, S1 and S5 Tables). Furthermore, we found that genes upregulated in *pals-17* mutants significantly correlate with genes that are differentially expressed (both upregulated and downregulated) in animals infected with other pathogens that are not associated with the IPR. *pals-17 pals-20* mutants upregulate genes in common with several other stress responses. In summary, we found that the loss of *pals-17* causes extensive changes in the transcriptional profile of animals, leading to the upregulation of the majority of IPR genes and other previously characterized immune and stress responses.

### PALS-17 and PALS-20 proteins are co-expressed in the intestinal cells

To examine where these newly identified regulators of the IPR are expressed, we created transcriptional and translational reporters for *pals-17* and *pals-20*. To first determine the tissues where they are expressed, we examined the expression of *pals-17*p::GFP and *pals-20*p::wrmScarlet transcriptional reporters at the L4 stage. Both transgenes were expressed in intestinal tissue, especially in the anterior-most and posterior-most intestinal cells (Fig 5A and 5B). Furthermore, we found that both reporters were expressed in the head region. *pals-17*p::GFP signal was present in head neurons (Fig 5A), whereas *pals-20*p::wrmScarlet was sporadically and weakly expressed in the head region (Fig 5B).

**Fig 5.**
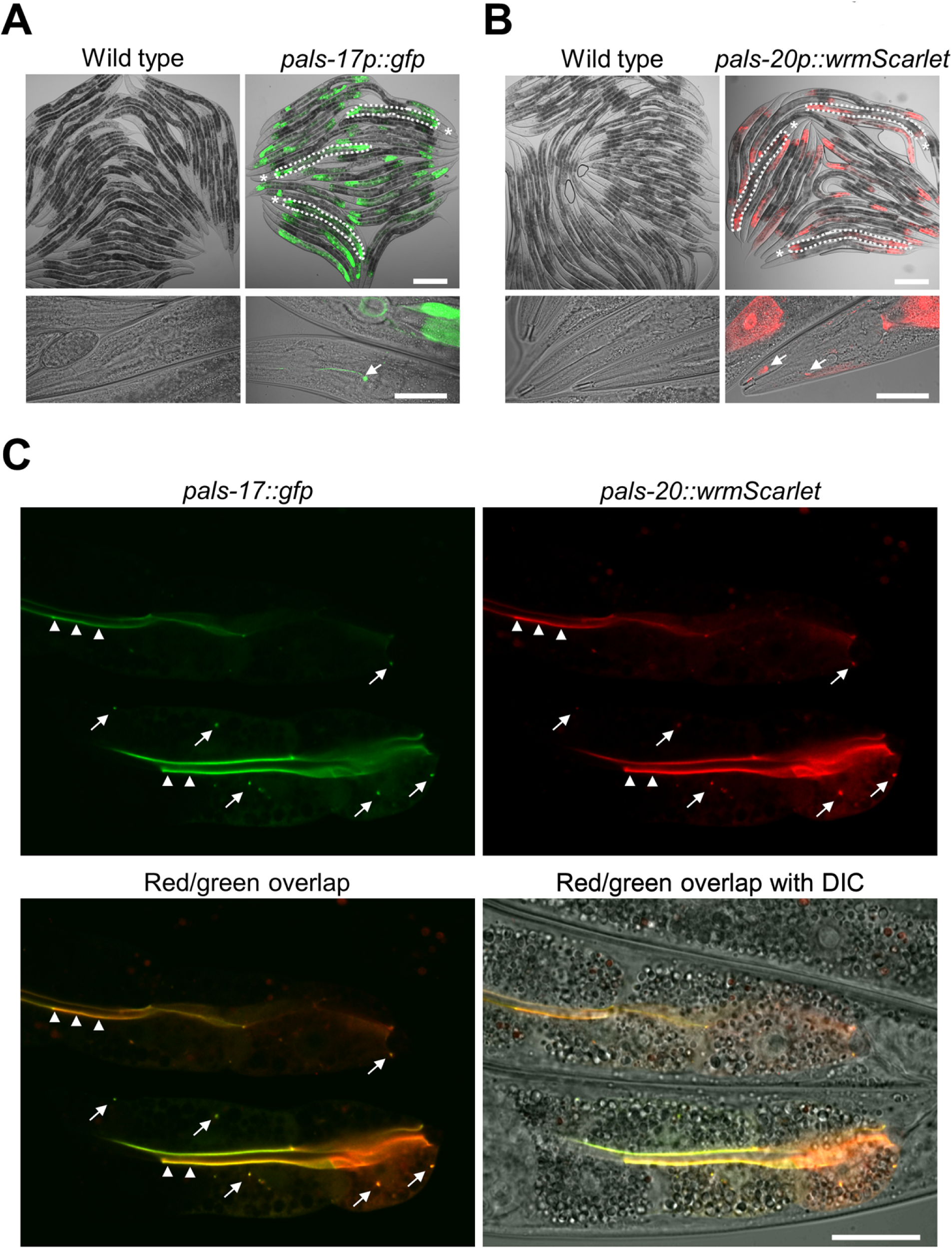
PALS-17 and PALS-20 colocalize in the intestine. (A, B) Expression of *pals-17*p::GFP (A) and *pals-20*p::wrmScarlet transcriptional reporters (B) in the intestine (indicated for a few worms with white dotted line in upper images; asterisks indicate heads) and the head region (indicated with white arrows in lower images). For comparison, the wild-type control is provided for each transgenic strain. Scale bars, 200 µm (upper images) and 50 µm (lower images). (C) Expression of PALS-17::GFP and PALS-20::wrmScarlet translational reporters and their colocalization. Representative images showing separate red and green channels, as well as their overlap and overlap with DIC, are shown. Arrowheads indicate colocalization of fluorophores at the border with the intestinal lumen; arrows indicate areas of colocalization in other regions of intestinal cells. Scale bar, 20 µm.

To examine if PALS-17 and PALS-20 proteins colocalize in intestinal cells, we co-expressed PALS-17::GFP and PALS-20::wrmScarlet translational reporters. We found that these two proteins formed puncta in a subset of intestinal cells, and in most cases, they colocalized with each other (Fig 5C). Furthermore, we observed fluorescent signals from both reporters at the apical border of the intestinal cells, lining the intestinal lumen. To confirm that these translational reporters are functional, we tested if the expression of PALS-17::GFP can suppress the growth defects of *pals-17* mutants and if PALS-20::wrmScarlet overexpression slows down the development of *pals-17 pals-20* double mutants. We found that both constructs are functional, because their expression had significant effects on the growth of animals carrying the array, as compared to their non-transgenic siblings (S6 Fig, S1 Table). Unfortunately, we were unable to amplify or synthesize the DNA region upstream of *pals-16* to obtain a *pals-16* reporter, likely because of the repetitive and complex sequence in this region. In summary, we found that PALS-17 and PALS-20 proteins colocalize in the intestine, which is the tissue invaded by viral and microsporidian pathogens that activate the IPR.

### Loss of *pals-17* increases resistance against intracellular pathogens in a *pals-20*-and *pals-16*-dependent manner

IPR induction in the *pals-22* mutant background leads to increased resistance against intestinal intracellular pathogens, namely the Orsay virus and the microsporidian species *N. parisii*. Therefore, we examined whether *pals-17* mutants show increased resistance against these pathogens, and if so, whether this resistance is *pals-20*- and/or *pals-16*-dependent. First, we analyzed Orsay virus infection levels after infecting L1 animals for 18 hours and found a significantly lower infection level in *pals-17* mutants (Fig 6A, S1 Table). Loss of either *pals-20* or *pals-16* suppressed this phenotype, suggesting that both genes contribute to the increased viral resistance of *pals-17* mutants. We also examined antiviral resistance by infecting L4 animals and measuring their infection levels at 24 hours post-inoculation (hpi). To increase the overall infection level, animals were subjected to *rde-1(RNAi)* from the L1 stage. *rde-1* plays an important role in antiviral immunity as a critical component of the RNAi pathway, but it does not have a known role in regulating the IPR (19). Here again, we saw that *pals-17* mutants showed significant resistance to Orsay virus infection, which was rescued by the loss of *pals-20* and/or *pals-16* (Fig 6B, S1 Table). This result indicated that *pals-20* is still required for immunity at later stages of life, although its role in IPR gene expression seems to be restricted to younger animals.

**Fig 6.**
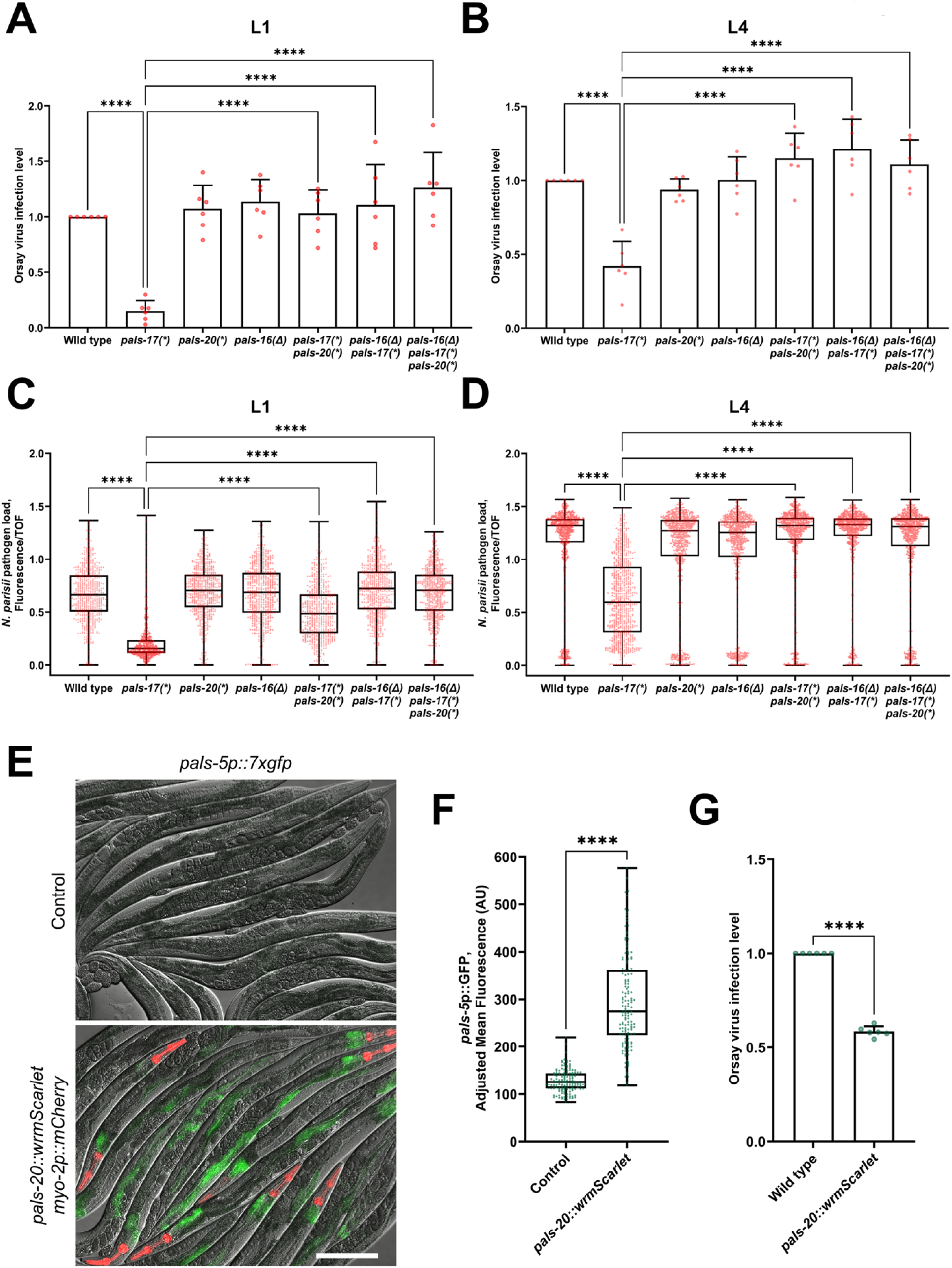
*pals-17*, *pals-20* and *pals-16* play a role in immunity against intracellular pathogens. (A, B) Fractions of animals infected with the Orsay virus in indicated genetic backgrounds. Animals were inoculated with the pathogen from the L1 (A) or L4 stage (B). Animals were scored based on the presence or absence of the Orsay virus-specific FISH probe fluorescence (three experimental replicates, at least 300 animals per replicate). Red dots represent the infection level for each replicate; the wild type was set at one for each replicate. Error bars are SD. An ordinary one-way ANOVA test was used to calculate p-values; **** p < 0.0001. (C, D) Quantification of *N. parisii*-specific FISH fluorescence signal standardized to time-of-flight (TOF). Animals were inoculated with pathogen from the L1 (C) or L4 stage (D), and analyzed at 30 hpi. 600 (C) and 750 animals (D) were analyzed per strain in three experimental replicates. Red dots in the box-and-whisker plots represent individual values for each analyzed animal. Box lines represent median values, box bounds indicate 25^th^ and 75^th^ percentiles, and whiskers extend to the minimum and maximum values. A Kruskal-Wallis test was used to calculate p-values; **** p < 0.0001. (E, F) *pals-5*p::7xGFP reporter expression is induced in animals overexpressing *pals-20::wrmScarlet* construct. (E) Representative images of synchronized animals 72 hours-after L1 overexpressing *pals-20::wrmScarlet myo-2p::mCherry* array and their siblings that do not overexpress *pals-20* (control). Red, green and DIC channels were merged. *myo-2*p::mCherry co-injection marker is shown in red. PALS-20::wrmScarlet is not visible with the exposure setting that was used for imaging. Scale bar, 200 µm. (F) Quantification of *pals-5*p::7xgfp expression in *pals-20::wrmScarlet* overexpressing strain and their siblings that that do not overexpress *pals-20* (control). Green dots in the box-and-whisker plots represent individual values for each analyzed animal; 50 animals were analyzed per strain for each of the three experimental replicates. Box lines represents median value, box bounds indicate 25^th^ and 75^th^ percentiles, and whiskers extend to the minimum and maximum values. A Kolmogorov-Smirnov test was used to calculate p-values; **** p < 0.0001. AU = arbitrary units. (G) Fractions of animals overexpressing *pals-20::wrmScarlet* construct and their non-transgenic siblings that are infected with Orsay virus. Animals were scored based on the presence or absence of the Orsay virus-specific FISH probe fluorescence (three experimental replicates, at least 300 animals per replicate). Green dots represent the infection level for each replicate; the wild type was set at one for each replicate. Error bars are SD. An unpaired t-test was used to calculate the p-value; **** p < 0.0001.

Next, we analyzed the resistance of *pals* mutants to the microsporidian species *N. parisii*. We fed animals *N. parisii* spores at either the L1 and L4 stages, and then examined pathogen load in the intestine at 30 hpi. At this time point *N. parisii* develop into multinucleate proliferative forms known as meronts. Here we observed significantly lower pathogen load in *pals-17* mutants at both analyzed stages (Fig 6C and 6D, S1 Table). The intestinal cells of *pals-17 pals-20* double mutants that were inoculated with pathogen from the L1 stage also had a significantly lower amount of meronts relative to their size in comparison to the wild-type control. However, the pathogen load of *pals-17 pals-20* double mutants was higher than in *pals-17* single mutants, indicating partial suppression of this phenotype by *pals-20* at the L1/L2 stage. Alternatively, the difference between these two genetic backgrounds could be a consequence of differences in developmental rates. Interestingly, *pals-17 pals-20* double mutants inoculated at the L4 stage had wild-type pathogen load levels, again suggesting that *pals-20* promotes resistance at later stages of life. A *pals-16* mutation fully suppressed the increased pathogen resistance observed in *pals-17* mutants at both time points.

In order to determine whether the reduced pathogen load of *pals-17* mutants is due to reduced feeding and, therefore, reduced exposure to pathogens in the lumen, we fed fluorescent beads to L1 animals and measured bead accumulation in the intestinal lumen. Here we found that *pals-17* mutants have significantly reduced fluorescent bead accumulation compared to wild-type animals (S7A Fig, S1 Table). Of note, *pals-17 pals-20* double mutants also showed significantly lower fluorescent bead accumulation, although these mutants had wild-type infection levels in many assays. This result suggests that feeding rate may not be a determining factor in these infection assays.

To address the concern that the decreased pathogen load in *pals-17* mutants, dependent on IPR activators like *pals-20,* was solely due to decreased exposure to pathogens, we examined the pathogen resistance and the feeding rate of a wild-type strain overexpressing a functional PALS-20::wrmScarlet construct. Importantly, unlike in the *pals-17 pals-20* double mutant background, we did not observe any effect of PALS-20::wrmScarlet overexpression on the developmental rate of wild-type animals, and these animals had normal rates of fluorescent bead accumulation (S7B Fig, S1 Table). First, we tested if PALS-20::wrmScarlet overexpression had an effect on IPR activation. We found high induction of IPR reporter expression in *pals-20* overexpressing animals in comparison to their wild-type siblings (Fig 6E and 6F, S1 Table). Next, we investigated Orsay virus infection levels in these two groups of animals and found that PALS-20::wrmScarlet overexpression leads to a significant reduction in Orsay virus infection level (Fig 6G, S1 Table). This result confirms the role of *pals-20* in controlling immunity against intracellular pathogens. In summary, loss of *pals-17* leads to significantly lower infection levels with both the Orsay virus and microsporidia. Loss of *pals-20* and *pals-16* suppress infection phenotypes of *pals-17* mutants in different stages of life, and overexpression of *pals-20* provides enhanced immunity against infection.

## DISCUSSION

In this study, we identified three previously uncharacterized genes *pals-17*, *pals-20*, and *pals-16* as important new regulators of the balance between immunity and development (Fig 7). The expanded *pals* gene family in *C. elegans* consists of two groups of genes: genes that are upregulated upon infection with obligate intracellular viral and fungal pathogens – “Induced *pals* genes”, and genes that do not change expression levels upon infection – “Uninduced *pals* genes”. *pals-17*, *pals-20*, and *pals-16* all belong to the latter group, together with the previously characterized genes *pals-22* and *pals-25*, which act as antagonistic regulators of the IPR and growth (S4 Fig) (4). Through a forward genetic screen, we identified *pals-17* as a negative regulator of the IPR. We demonstrate that *pals-17* is essential for normal development, as deletion of *pals-17* leads to larval arrest. Our RNAi suppressor studies identified two other *pals* family members, *pals-20* and *pals-16*, which function as *pals-17* antagonists by promoting IPR activation and inhibiting normal larval development. The negative regulator protein PALS-17 and the positive regulator protein PALS-20 colocalize in intestinal epithelial cells in discrete cytoplasmic puncta, as well as along the apical plasma membrane. Finally, we demonstrate that loss of *pals-17* as well as overexpression of *pals-20* protect animals against infection by intracellular pathogens. Because previous studies have only functionally characterized two out of 39 *pals* genes, our findings have more than doubled the number of *pals* genes with known functions.

**Fig 7.**
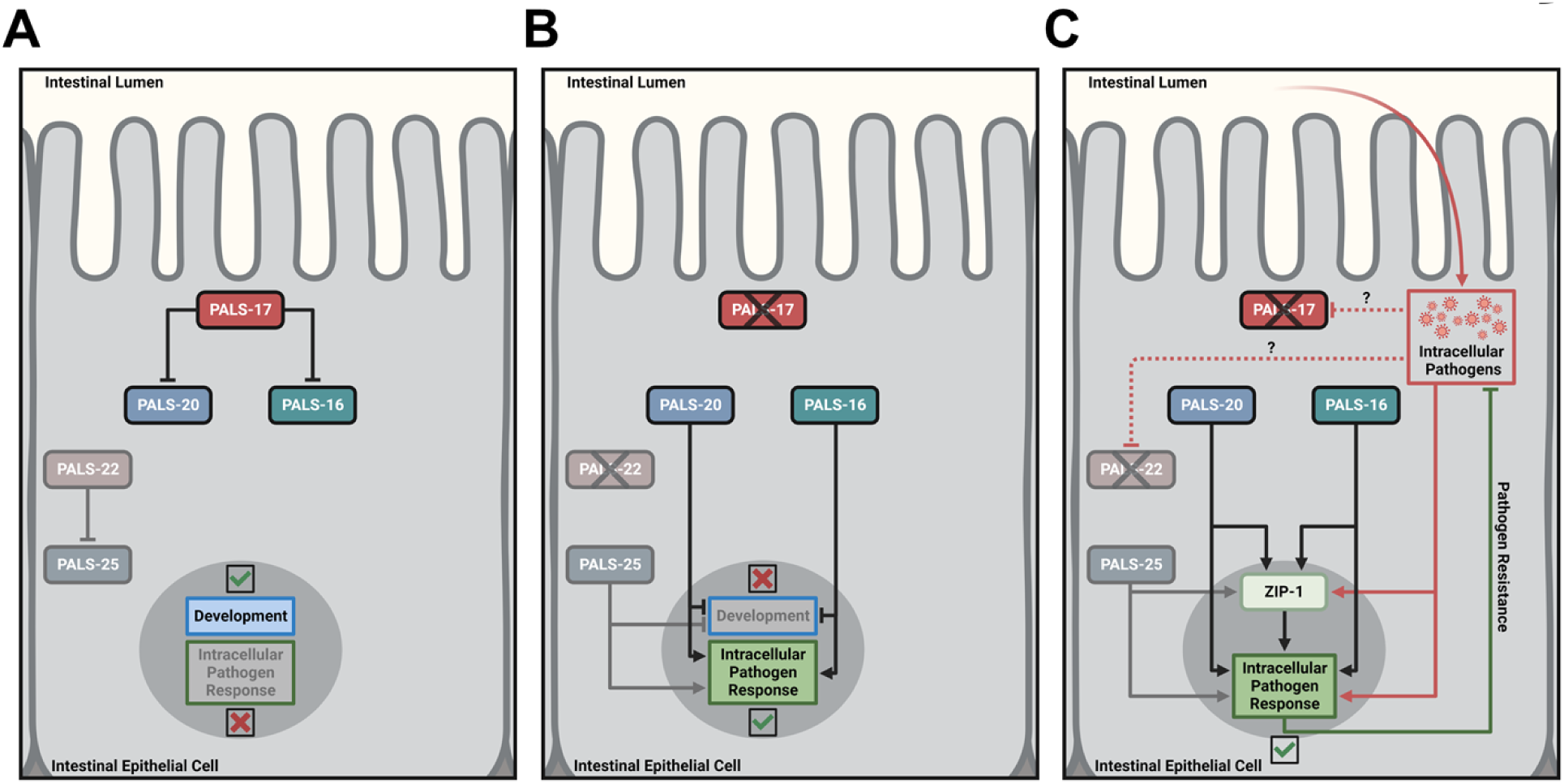
Models for regulation of development and immunity by *pals-17*, *pals-20* and *pals-16*. (A) PALS-17 is a negative IPR regulator acting upstream of the positive IPR regulators PALS-20 and PALS-16, similar to how PALS-22 is a negative IPR regulator acting upstream of the positive IPR regulator PALS-25. (B, C) In the absence of PALS-17, PALS-20 and PALS-16 inhibit development and activate the IPR in *C. elegans*, similar to PALS-25. Created with BioRender.com.

Strong selective pressures imposed by pathogens are likely the main drivers of the evolutionary expansion of the *pals* gene family in *C. elegans*. Such examples of species-specific immune gene family expansions are numerous in eukaryotes (12). For example, the family of immunity-related GTPase encoding genes in mice likely expanded as a consequence of the arms race with rhoptry kinases from the protozoan parasite *Toxoplasma gondii*, as well as with other intracellular pathogens including microsporidia (12, 20-22). Furthermore, the plant immunity R genes are expanded in plant genomes and share many characteristics in common with the *C. elegans pals* gene family, although they are not homologs (4). In particular, there are similarities in gene expression regulation, protein-protein interactions and phenotypes between the *pals-22/pals-25* pair and the R gene pair RPS4/RRS1 in the plant *Arabidopsis thaliana* (23-26). Importantly, R genes regulate a switch between growth and defense, similar to *pals* genes (1-4).

The presence of antagonistic paralogs in *pals* and R gene families highlights the importance of maintaining the balance between immunity and development. In this study, we demonstrate the striking effect that loss of *pals-17* has on animal development, which can be reversed by loss of either *pals-20* or *pals-16*. Interestingly, mutations in *pals-22* also cause developmental delay, but to a lesser degree than the loss of *pals-17* (4-6). Although *pals-22(jy1)* and *pals-17(*)* alleles have premature stop codons in similar regions of *pals-22* and *pals-17*, developmental impairment is more severe in *pals-17(*)* animals. Furthermore, complete loss of *pals-17* causes early developmental arrest, implying that the severity of developmental phenotypes in different *pals-17* mutants may depend on the expression levels of *pals-17*-dependent genes, including IPR genes.

Other immunity-related genes have been previously implicated in organismal development. For example, overexpression of immune genes, which participate in defense against the bacterial pathogen *Pseudomonas aeruginosa*, have been shown to impair development of *C. elegans*. Specifically, hyperactivation of PMK-1 p38 MAPK pathway components in animals that are not exposed to pathogen causes significant delay in development (27, 28). Recently, gut microbiota have been shown to regulate activation of this immune pathway, affecting organismal growth on certain food sources (29). Furthermore, a null mutation in the gene encoding pseudokinase NIPI-3, an important immunoregulator, leads to developmental arrest (30, 31). NIPI-3 is a negative regulator of bZIP transcription factor CEBP-1, which stimulates transcription of *sek-1* – a core component of the PMK-1 p38 MAPK immune pathway during development (28, 31, 32). Loss-of-function mutations in several genes from the PMK-1 p38 MAPK pathway suppress the larval arrest phenotype of *nipi-3* mutants, suggesting that immune hyperactivation has a detrimental effect on development (28). Similarly, overexpression of a cytokine, tumor necrosis factor (TNF), causes growth impairment in mice, due to improper cartilage development and insufficient bone growth (33). The elevated levels of different cytokines also play important roles in the intrauterine development of mammals. For example, increased levels of interferon-gamma (IFN-γ), type-I IFNs, TNF-α, and interleukins 1β, 6 and 17 lead to developmental failures in different fetal organs, including the neural tube, brain, lungs and placenta (34-42). These examples demonstrate the deleterious effects that an overstimulated immune system can have on growth and reproduction.

The balance between immunity and development is also dependent on the timing of gene expression. Our data demonstrate that IPR gene upregulation in *pals-17* mutants can be completely suppressed by loss of *pals-16* at different developmental stages, whereas loss of *pals-20* specifically suppresses this phenotype in L1 larvae and not in L4 larvae. Nevertheless, we observed that *pals-20* still plays a role in immunity at later stages of development. This result suggests that IPR activation in L1 animals could be essential for priming animals for defense against pathogens that they may encounter later in life. Therefore, the lack of IPR induction in L1 *pals-17 pals-20* mutants might be sufficient to cause wild-type susceptibility to intracellular pathogens later in life. Alternatively, it is possible that IPR activation by *pals-20* fluctuates throughout life and may not be specific for younger developmental stages. Furthermore, our RNA-seq analyses show a drastic difference in the number of differentially expressed genes between partially synchronized *pals-17* and L4 stage *pals-17 pals-20* mutant animals. This distinction suggests that there may be some other *pals-17*-dependent genes that play a role in immunity and are still suppressed by loss of *pals-20* at the L4 stage.

Altogether, in this and our previous studies, we have shown that five uninduced *pals* genes, *pals-16, pals-17, pals-20, pals-22 and pals-25*, act as ON/OFF switch modules to regulate induced *pals* genes and other IPR genes (4-6). Our analysis indicates that the “OFF switch” *pals-17* likely functions in parallel to the “OFF switch” *pals-22*. Predicted protein structures of regulatory PALS proteins (S4 Fig) suggest that similar secondary protein structure corresponds to similar function. It is possible that other uninduced *pals* genes function in a similar manner to the *pals-16/pals-17/pals-20* module and the *pals-22/pals-25* module. Further studies could address if other closely related uninduced *pals* genes represent additional modules that regulate the IPR and development. Having several parallel modules that can be interchangeably activated likely provides an advantage for hosts defending against infection. Such modules exist among plant R genes, where the same bacterial effector AvrRps4 from *Pseudomonas syringae* can be recognized by two parallel R gene modules – RPS4/RRS1 and RPS4B/RRS1B (25). However, in other contexts, R genes act as specialized sensors that recognize distinct effectors from plant pathogens. For example, the PopP2 effector from bacterial species *Ralstonia solanacearum* is specifically recognized by the RPS4/RRS1 gene pair, but not the RPS4B/RRS1B gene pair (25). While it is not known what activates *pals* genes in *C. elegans*, is possible that different *pals* modules in *C. elegans* are responsible for sensing different pathogen effectors in order to induce the IPR. If so, it would help provide an explanation for the expansion of the *pals* gene family in *C. elegans*, which appears to have extensive control over the balance between immunity and development.

## MATERIALS AND METHODS

### *C. elegans* maintenance

*C. elegans* strains were grown on Nematode Growth Media (NGM) agar plates with streptomycin-resistant *Escherichia coli* OP50-1 at 20 °C (43), unless stated otherwise. Strains used in this study are listed in S6 Table.

### Synchronization of *C. elegans*

To obtain synchronized populations of *C. elegans* (44), gravid adult animals were washed off plates with M9 media and collected into 15 ml conical tubes. Tubes were centrifuged and the supernatant was removed leaving 3 ml of M9. 1 ml of bleaching solution (500 μl of 5.65–6% sodium hypochlorite solution and 500 μl of 2 M NaOH) was then added to the tube until adult animals were partially dissolved. M9 was added to 15 ml to wash the released embryos. The tubes were immediately centrifuged and the supernatant was removed. The washing step was repeated four times and embryos were resuspended in a final volume of 3 ml of M9. Embryos were placed in a 20 °C incubator with continual rotation for 20–24 h to hatch into L1 animals.

### Chemical mutagenesis, forward genetic screen and mapping of *pals-17(jy74)*

L4 stage *pals-22(jy1) pals-25(jy11)* mutants carrying the *jyIs8[pals-5p::gfp myo-2p::mCherry]* transgene were washed off plates with M9 media and collected into 15 ml conical tubes. Following centrifugation and supernatant removal, animals were washed with M9 two times to remove bacteria. The worm pellet was resuspended in 4 ml M9 and 25 µl of ethyl methanesulfonate (EMS) was added to the tube (45). Animals were incubated at room temperature for 4 h with continual rotation. After incubation, the worms were pelleted and the supernatant was removed. Animals were washed with M9 two times to remove the mutagen. Worms were plated on 10 cm Petri plates with OP50-1 bacteria and incubated at room temperature for 2.5 h to recover. 18 groups of 30 mutagenized animals were then transferred to new 10 cm Petri plates with OP50-1 bacteria and incubated at 20°C. After 36 h, mutagenized P0 animals were removed and the incubation of F1 progeny was continued. 16,752 F1 animals reached adulthood after three days and they were bleached to obtain their synchronized L1 progeny (thus, we calculate that we screened 33,504 haploid genomes). These F2 progeny were grown and analyzed for expression of *pals-5*p::GFP at different developmental stages, which led to the isolation of *jy74* mutants. These mutants were monitored through several generations to confirm that the GFP ON phenotype was still present. *jy74* animals were backcrossed to wild-type strain six times.

Genomic DNA from wild-type and mutant strains was isolated using Qiagen Puregene Core Kit A and submitted to BGI Genomics for whole genome sequencing. The Galaxy platform (https://usegalaxy.org) was used for mapping and variant calling to identify the causative mutation in the *jy74* allele. The quality check of the reads was performed using FastQC. Bowtie2 program (46, 47) was used to map the reads against the reference *C. elegans* genome WBCel235.dna.toplevel.fa.gz. Genetic variants were identified using the FreeBayes program. The VcfAllelicPrimitives program was used to split allelic primitives into multiple VCF lines. Wild-type and mutant VCF datasets were intersected using the VCF-VCFintersect program. Variants were filtered using the filter criteria (QUAL>30)&(DP>=10)&(isHOM(Gen[0]) in SnpSift Filter (48), and the remaining variants were annotated using the SnpEff eff program (49).

### Imaging

Before imaging, animals were anesthetized with 100 µM levamisole. Samples were imaged using a Zeiss AxioImager M1 compound microscope equipped with Axio Vision 4.8.2 software (Fig 1A, 1B, 1F, 1G, 1I, 2A, 3A, 6E, S1, S2B and S3A) or a Zeiss LSM700 confocal microscope run by ZEN2010 software (Fig 5).

### *pals-5*p::GFP fluorescence measurements

Following 24 h or 44 h incubation at 20 °C, synchronized animals were collected and washed in M9 and loaded into 96-well plates. GFP fluorescence and time-of-flight (TOF) were measured using the COPAS Biosort machine (Union Biometrica). Fluorescence intensity was standardized to the TOF values, which are a proxy for worm length. Data analysis was performed using GraphPad Prism 9. Raw data are shown in S1 Table. For measurements of the *pals-5*p::7xGFP reporter expression, animals were first fixed in 4% paraformaldehyde for 5 min and then imaged using the ImageXpress automated imaging system NanoImager. Fluorescence was measured using FIJI software (50) as a mean fluorescence per animal, which was normalized to background florescence.

### Body length measurements

Synchronized animals were grown at 20 °C for 44 h, collected and washed in M9, and fixed in 4% paraformaldehyde for 5 min. Animals were imaged using the ImageXpress automated imaging system NanoImager (Molecular Devices, LLC) in 96-well plates. The length of each animal was measured using the FIJI program (50). Fifty animals were analyzed for each strain, for each of the three experimental replicates. Data analysis was performed using GraphPad Prism 9.

### RNAi assays

RNA interference assays were performed using the feeding method (51). Overnight cultures of HT115 *E. coli* were plated on RNAi plates (NGM plates supplemented with 5 mM IPTG and 1 mM carbenicillin) and incubated at room temperature for 1-3 days. Synchronized L1 or L4 animals were transferred to these plates and incubated at 20 °C. The effects of *pals-17* RNAi treatment (Fig 1G and 1I) on animals that were plated at the L1 stage were examined after 44 h incubation. For *pals-20* RNAi treatment (Fig 2A), the progeny of L4 animals were examined after three days. The effects of *pals-16* RNAi (Fig 3A) were examined 72 h after plating L1 animals. Control RNAi experiments were carried out using vector-alone plasmid L4440.

### Generation of CRISPR/Cas9-mediated mutations in *pals-17*, *pals-20* and *pals-16* genes

We designed mutations for *pals-17* alleles *syb4005* (*pals-17(Δ)*) and *syb3980* (*pals-17(*)*) and *pals-20* allele *syb3744* (*pals-20(*)*); CRISPR/Cas9-mediated genome editing was performed by SunyBiotech. SunyBiotech balanced *pals-17(Δ)* allele with *sC1(s2023)*. *pals-17(Δ)* allele is a 1,394 base pair deletion and a 16 base pair insertion. The deletion starts 36 nucleotides upstream of the *pals-17* start codon and ends at the 69^th^ nucleotide after the stop codon; the inserted sequence is 5’-AAAGATCTTTATTTAA-3’. The *pals-17(*)* allele contains six point mutations that change amino acids L(49) N(50) and Y(51) to three consecutive stop codons; sequence 5’-CTTAACTAC-3’ is mutated to 5’-TAGTAATAA-3’, creating a SpeI cutting site. *pals-20(*)* allele contains two point mutations in *pals-20* that change amino acid E(15) to stop codon and one silent mutation in the 13^th^ codon encoding valine; sequence 5’-GTTATCGAA-3’ is mutated to 5’-GTCATCTAG-3’, creating a XbaI cutting site. The mutations in *pals-20(*)* should spare the function of the *Y82E9BR.5* gene, which completely overlaps with *pals-20.* Genotyping primers for *pals-17* (primers 1 and 2) and *pals-20* point mutants (primers 3 and 4) are listed in S7 Table.

The *pals-17* allele *jy167* was created in the *pals-20* mutant background; the *jy167* allele contains the same mutations as *syb3980* and they are both labeled as *pals-17(*)* in the main text. The deletion alleles of *pals-16* in the wild-type, *pals-17*, and *pals-17 pals-20* mutant backgrounds are *jy182*, *jy171* and *jy174*, respectively; all three alleles are labeled as *pals-16(Δ)* throughout the text. The *jy167*, *jy182*, *jy171* and *jy174* alleles were all created using the co-CRISPR method with preassembled ribonucleoproteins (52, 53). Cas9-NLS protein (27 µM final concentration) was obtained from QB3 Berkeley; crRNA, tracrRNA and DNA primers were ordered from Integrated DNA Technologies (IDT). The crRNA sequence used for generating the *jy167* allele in *pals-17* was ATTCAGGTTCCATGTAGTTA; the repair template sequence was 5’-TGGAGAGGCAAGGCGATTTTGCTAGACCAGAGCAGTCAAACTAGTAATAAATGGAACCTGAATATTATGAGTTTAAGGTATATTTTATTTTGAGAATATATAA-3’. This repair template sequence replaces wild-type sequence 5’-CTTAACTAC-3’ with mutated sequence 5’-TAGTAATAA-3’ that changes amino acids L(49) N(50) and Y(51) to three consecutive stop codons and adds a SpeI cutting sit. The mutated region is flanked with a 41-nucleotide overhang on the 5’ end and a 53-nucleotide overhang on the 3’ end. AAACGTTTCTACACAGCATG and ACATCGTGACGCCAAATTCA crRNA sequences were used to target regions of the *pals-16* gene close to the start codon and the stop codon, respectively. The *jy182* allele is a 1,300 base pair deletion in *pals-16* in the wild-type background, starting one nucleotide upstream of the *pals-16* start codon and ending 14 nucleotides upstream of the stop codon. The *jy171* allele is a 1299 base pair deletion and seven base pair insertion in *pals-16* in the *pals-17(*)* background. The deletion starts at the second nucleotide of the *pals-16* start codon and it ends 13 nucleotides upstream of the stop codon; the inserted sequence is 5’-CGGGTGA-3’. The *jy174* allele is a 1,302 base pair deletion in *pals-16* in the *pals-17(*) pals-20(*)* background, starting immediately after the *pals-16* start codon and ending seven nucleotides upstream of the stop codon. PCR genotyping was performed using primers 5-7 listed in S7 Table. Selected lines were backcrossed three times to the wild-type strain before they were used in experiments.

### Developmental rate assays

Gravid adult animals were grown at 20 °C and picked onto 10-cm seeded NGM plates (40 animals per strain; 1 plate per strain). Due to the low brood size of *pals-17* mutants, 60 animals were picked for this strain to ensure an adequate sample size of progeny. Animals were then incubated at 20 °C for 2 h. Next, approximately 7 mL of M9 was poured onto the plate and gently swirled for 15-20 s to dislodge all adult animals while leaving eggs fastened to the *E. coli* OP50-1 lawn. The M9 was removed, and the plates were thoroughly checked under a microscope to ensure that no adult animals remained. After verifying that the eggs were not washed off, the plates were incubated at 20 °C and the time course began. At the 44 h, 64 h, and 72 h time points, 100 animals on each plate were scored for development as being at or below the L3 stage or at or above the L4 stage. Data analysis was performed using GraphPad Prism 9. Raw data are shown in S1 Table.

### RNA isolation

Total RNA isolation was performed as previously described (54). For RNA isolation from L1 animals, 35,000 worms were grown per NGM plate for 3 h at 20 °C; one or two plates were used per strain per replicate. For RNA isolation from L2 animals, 10,000 worms were grown per NGM plate for 33 h (*pals-17(*)* mutants) or 24 h (all other strains) at 20 °C; four plates were used per strain per replicate. For RNA isolation from L4 animals, 3,000 worms were grown per NGM plate for 72 h (*pals-17(*)* mutants) or 48 h (all other strains) at 20 °C; up to four plates were used per strain per replicate. Animals were washed off plates using M9, then washed with M9 and collected in TRI reagent (Molecular Research Center, Inc.). RNA was isolated using a BCP phase separation reagent, followed by isopropanol and ethanol washes. For RNA-seq analysis, samples were additionally purified using the RNeasy Mini kit (Qiagen).

### Quantitative RT-PCR

qRT-PCR analysis was performed as previously described (54). In brief, cDNA was synthesized from total RNA using iScript cDNA synthesis kit (Bio-Rad). qRT-PCR was performed using iQ SYBR Green Supermix (Bio-Rad) with the CFX Connect Real-Time PCR Detection System (Bio-Rad). Three independent experimental replicates were performed for each qRT-PCR analysis. Each sample was analyzed in technical duplicates. All values were normalized to the expression of the *snb-1* control gene, which does not change expression upon IPR activation (6). The Pffafl method was used for data quantification (55). The sequences of the primers used in all qRT-PCR experiments are given in S7 Table (primers 8-25). Raw data are shown in S1 Table.

### Phylogenetic analysis of PALS proteins

Amino acid sequences were obtained from https://wormbase.org/. 38 out of 39 PALS proteins were analyzed; PALS-36 was excluded from the analysis because it is annotated as a pseudogene and the transcript is not available. Eight PALS proteins have more than one isoform: isoform “a” transcripts were used for PALS-3, PALS-7, PALS-11, PALS-19, PALS-23, PALS-37 and PALS-38; the isoform “b” transcript was used for PALS-31. Clustal omega program (https://www.ebi.ac.uk/Tools/msa/clustalo/) was used for sequence alignment of 38 PALS protein sequences (56). A phylogeny software PhyML 3.0 (http://www.atgc-montpellier.fr/phyml/) based on the maximum likelihood principle with the Smart Model Selection criterion (57, 58) was used to create a radial phylogram shown in S4A Fig. The Sequence Manipulation Suite: Color Align Conservation program (https://www.bioinformatics.org/sms2/color_align_cons.html) was used for visualization of amino acid sequence alignment between PALS-20, PALS-16 and PALS-17 in S4B Fig. Black boxes represent identical amino acids; grey boxes represent similar amino acids as defined by seven similarity groups: GAVLI, FYW, CM, ST, KRH, DENQ, P. Percent identity and similarity between protein sequences shown in S2 Table were calculated using the Sequence Manipulation Suite: Ident and Sim program (https://www.bioinformatics.org/sms2/ident_sim.html).

### Protein structure models

Predicted structures of PALS proteins were obtained from AlphaFold Protein Structure Database (https://alphafold.ebi.ac.uk/) (15). The PyMOL Molecular Graphics System version 2.5.2, Schrodinger, LLC software was used for protein structure examination and overlap.

### RNA-sequencing analyses

cDNA library preparation and paired-end sequencing were performed at the Institute for Genomic Medicine at the University of California, San Diego. Reads were mapped to *C. elegans* WS235 genome using Rsubread in RStudio. Differential expression analyses were performed using limma-voom function in the Galaxy platform (https://usegalaxy.org/). Genes with counts number lower than one count per million (CPM) were filtered out. Differentially expressed genes had adjusted p-values lower than 0.05 (S3 Table). Overlaps between datasets shown in Venn diagrams (Fig 4A-C) were calculated using Multiple List Comparator software (https://molbiotools.com/listcompare.php). The significance of overlaps between datasets was calculated using Overlap Stats software (http://nemates.org/MA/progs/overlap_stats.html). The default number of genes (17,611) was used for statistical analyses.

### Analysis of enriched gene categories

Annotations of gene categories upregulated in *pals-17(*)* but not in the wild-type background (Fig 4D) and in *pals-17(*) pals-20(*)* but not in wild-type samples (Fig 4E) were performed using the online tool WormCat (http://www.wormcat.com/) (16). Data were visualized using GraphPad Prism 9 software. Raw data are shown in S1 Table.

### Brood size quantification

Animals were grown at 20 °C and then single-picked onto separate 3.5 cm tissue culture-treated NGM plates upon reaching the L4 stage. Every 24 h, the number of eggs and progeny on each plate was recorded and the gravid adult animal was transferred to a new plate to be counted the next day. This process was repeated daily until all animals laid no eggs for three days, at which point the counts were summed to calculate the total progeny for each individual. Three experimental replicates were performed. Data analysis was performed using GraphPad Prism 9. Raw data are shown in S1 Table.

### Functional expression analysis

Gene Set Enrichment Analysis (GSEA) software v4.2.3 was used for functional analysis of *pals-17(*)* and *pals-17(*)pals-20(*)* compared to wild type (Fig 4F) using the GSEAPreranked module (17, 18). Differentially expressed genes from RNA-seq expression data for *pals-17(*)* versus wild type and *pals-17(*) pals-20(*)* versus wild type were ranked based on Log2 fold changes and converted into a GSEA-compatible file. 93 previously published gene sets were used for comparison (4, 5, 7, 14, 19, 54, 59-77). For both analyses, a signal-to-noise metric of 1000 permutations was used with “no collapse” as gene symbols were pre-defined in the ranked gene list. Gene sets that had fewer than 15 genes represented in the RNA-seq data from this study were excluded from statistical analysis. Normalized Enrichment Scores from GSEAPreranked analysis described above were calculated with default parameters. A heatmap of the GSEA analysis was made using GraphPad Prism 9 software. Raw data are shown in S1 Table; full datasets are shown in S5 Table.

### Molecular cloning and transgenesis

*pals-17::gfp* and *pals-20::wrmScarlet* translational reporter constructs were assembled using NEBuilder HiFi kit (New England Biolabs) by fusing PCR products and gene fragments. The pET755 plasmid was used for Q5 polymerase-mediated (New England Biolabs) PCR amplification of the backbones that contain an ampicillin resistance cassette and *unc-54* 3’utr, and, in the case of *pals-20* reporters, a wrmScarlet::3xHA sequence (primers 26-28 in S7 Table). Promoter regions were PCR amplified using Q5 polymerase (New England Biolabs) from *C. elegans* genomic DNA using primers 29-32 listed in S7 Table. Amplified promoter regions contain 1,143 base pairs upstream of the *pals-17* start codon and 1,000 base pairs upstream of the *pals-20* start codon. *pals-17(cDNA)::gfp::3xHA* and *pals-20* cDNA sequences were synthesized as gene fragments by Twist Biosciences.

*pals-17p::gfp* and *pals-20p::wrmScarlet* transcriptional reporter constructs were assembled by PCR amplification of regions of interest from the previously created translational reporter constructs (*pals-17* and *pals-20* cDNA segments were excluded), which were subsequently circularized using NEBuilder HiFi kit (New England Biolabs). Primers 33-36 were used for creating *pals-17* and *pals-20* transcriptional reporter constructs (S7 Table).

The resulting plasmids were verified by restriction digest and sequencing across the insert. Transcriptional reporters were individually injected at concentrations of 50 ng/µl. Translational reporters (c = 13 ng/µl for each) were co-injected with the *gcy-8p::gfp* neuronal marker (c = 20 ng/µl) into a wild-type genetic background. For functionality testing, *pals-5*p::GFP expression analysis and Orsay virus infection assay, the *pals-17::gfp* and *pals-20::wrmScarlet* construct were separately injected at a higher concentration of c = 95 ng/µl, together with a *myo-2p::mCherry* pharyngeal marker (c = 5 ng/µl).

The *pals-5p::7xgfp* reporter (allele *knuSi886*) was created by InVivo Biosystems as a split-GFP system. Using the MosSCI transgenesis method, the *pals-5* promoter was used to drive expression of seven copies of the 11^th^ segment of the *gfp* gene, the *eft-3* promoter was used to drive expression of the remaining ten segments of *gfp* gene, as well as *unc-119* rescuing cassette, were inserted into the *ttTi5605* site on the second chromosome in the *unc-119* mutant background.

### Orsay virus infections

Orsay viral preps were prepared as previously described (7). Synchronized L1 animals were exposed to a mixture of OP50-1 bacteria and Orsay virus for 18 h at 20 °C (Fig 6A). For infection at the L4 stage (Fig 6B), synchronized animals were grown on *rde-1* RNAi plates (1,000 animals per plate, 2 plates per strain) for 72 h (*pals-17(*)* mutants) or 48 h (all other strains) at 20 °C. *rde-1* RNAi increases susceptibility to Orsay virus infection (78, 79). L4 animals were top-plated with a mixture of Orsay virus, OP50-1 and M9, and incubated at 20 °C for 24 h. Following incubation, animals were collected and washed in M9, and subsequently fixed in 4% paraformaldehyde for 30 min. Fixed worms were washed and then stained at 46 °C overnight using FISH probes conjugated to the red Cal Fluor 610 fluorophore (Biosearch Technologies), targeting Orsay virus RNA1 and RNA2 (80, 81). Analyses were performed visually using a Zeiss AxioImager M1 compound microscope; at least 300 animals were scored for the presence of the FISH fluorescent signal per strain per replicate. Due to relatively high variation in infection levels between replicates, the percentage of infected wild-type animals was normalized to one. Data analysis was performed using GraphPad Prism 9 software. Normalized data are shown in S1 Table.

### Microsporidia infections

*N. parisii* spores were prepared as previously described (82). Spores (1 million per plate) were mixed with food and L1 (Fig 6C) or L4 (Fig 6D) synchronized animals. To reach the L4 stage, worms were grown on NGM plates for 72 h (*pals-17(*)* mutants) or 48 h (all other strains) at 20 °C. During infection, animals were incubated at 25 °C for 30 h. Animals were collected and fixed in 4% paraformaldehyde for 30 min depending on the assay. Fixed worms were stained at 46 °C overnight using FISH probes conjugated to the red Cal Fluor 610 fluorophore (Biosearch Technologies), targeting microsporidia ribosomal RNA (81, 82). *N. parisii* pathogen load was measured with the COPAS Biosort machine (Union Biometrica). Data analysis was performed using GraphPad Prism 9. Raw data are shown in S1 Table.

### Bead feeding assays

1,200 synchronized L1 worms were mixed with 6 μl fluorescent beads (Fluoresbrite Polychromatic Red Microspheres, Polysciences Inc.), 25 μl 10X concentrated OP50-1 *E. coli* and M9 (total volume 300 ul) (5, 14, 54). This mixture was plated on 6 cm NGM plates, allowed to dry for 5 min and then incubated at 25 °C. After 5 min incubation, plates were transferred to ice, washed with ice-cold PBST and fixed in 4% paraformaldehyde for 30 min. Samples were washed with PBST and animals were imaged using the ImageXpress Nano Automated Imaging System (Molecular Devices, LLC). Fluorescence was analyzed in FIJI program; red fluorescence was analyzed in Fig S7A and green fluorescence was analyzed in Fig S7B. Fifty animals were analyzed for each strain, for each of the three experimental replicates. Data analysis was performed using GraphPad Prism 9. Raw data are shown in S1 Table.

## ACKNOWLEDGMENTS

This work was supported by NIH under R01 AG052622 and GM114139 to E.R.T., and the American Heart Association postdoctoral award 19POST34460023 to V.L. We thank Spencer Gang, Isabel Mejia, Lakshmi Batachari, Manish Grover, Michalis Barkoulas and Mario Bardan Sarmiento for helpful comments on the manuscript. We thank David Wang and Spencer Gang for their contributions in generating the *knuSi884* allele through InVivo Biosystems. We thank Yishi Jin for sharing the fosmid library. This publication includes data generated at the UC San Diego IGM Genomics Center utilizing an Illumina NovaSeq 6000 that was purchased with funding from a National Institutes of Health SIG grant (#S10 OD026929). The models in Fig 7 were created using BioRender.com.

## SUPPORTING INFORMATION

**S1 Fig.**
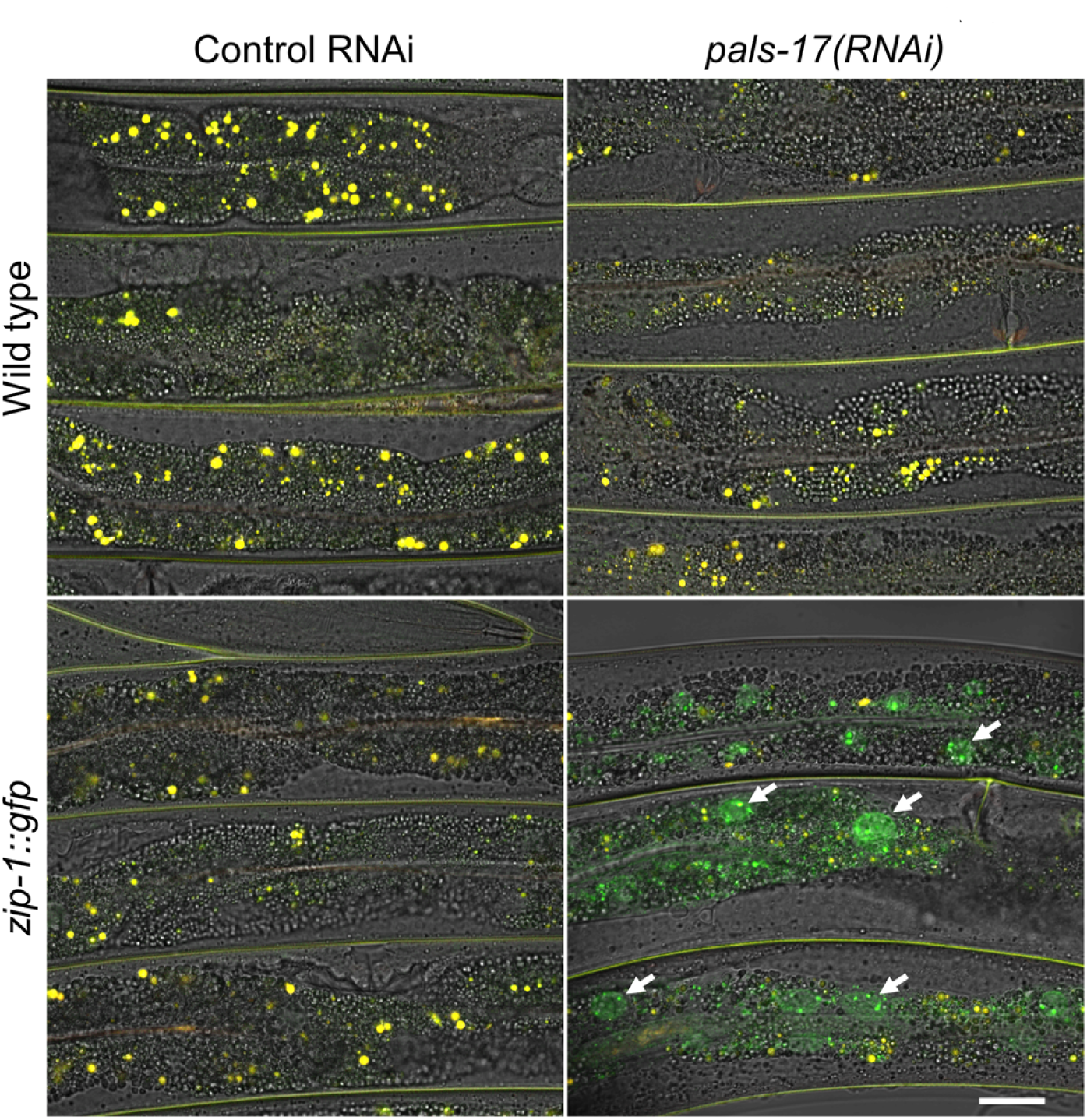
*pals-17(RNAi)* animals express ZIP-1::GFP in intestinal nuclei. Wild-type and *zip-1::gfp* animals treated with control and *pals-17* RNAi. Green, autofluorescence and DIC channels were merged. Intestinal ZIP-1::GFP expression is indicated with white arrows; autofluorescence from the gut granules and from the cuticle are shown in yellow. Scale bar, 20 µm.

**S2 Fig.**
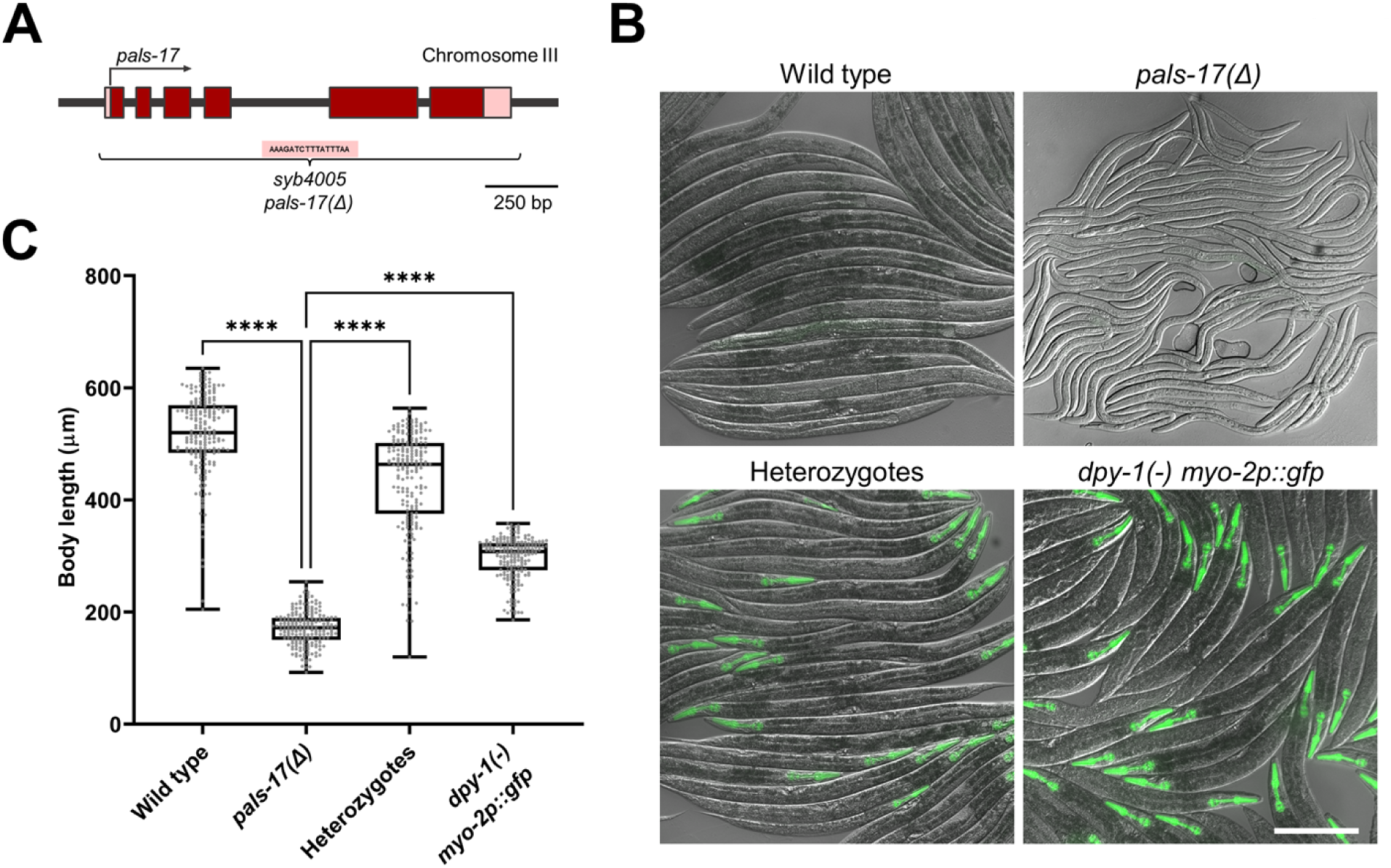
Deletion of *pals-17* causes developmental arrest. (A) *pals-17* gene structure. Exons are indicated with dark red boxes, 5’ and 3’ UTRs shown with light red boxes. The deletion in *syb4005* allele (*pals-17(Δ)*) is indicated by the bracket; the inserted sequence is shown in the pink rectangle. The horizontal arrow indicates the direction of transcription. (B) Synchronized *pals-17(Δ)* mutants and control strains following 44 h incubation at 20 °C. Green and DIC channels were merged. *myo-2*p::GFP present in the balancer strain is shown in green. Scale bar, 200 µm. (C) Box-and-whisker plot of body length values for indicated worm strains. Heterozygotes are *pals-17(Δ)*/*sC1(s2023)*. Box lines represent median values, box bounds indicate 25^th^ and 75^th^ percentiles, and whiskers extend to the minimum and maximum values. Gray dots represent individual values for each animal; 50 animals per each of the three replicates were analyzed. A Kruskal-Wallis test was used to calculate p-values; **** p < 0.0001.

**S3 Fig.**
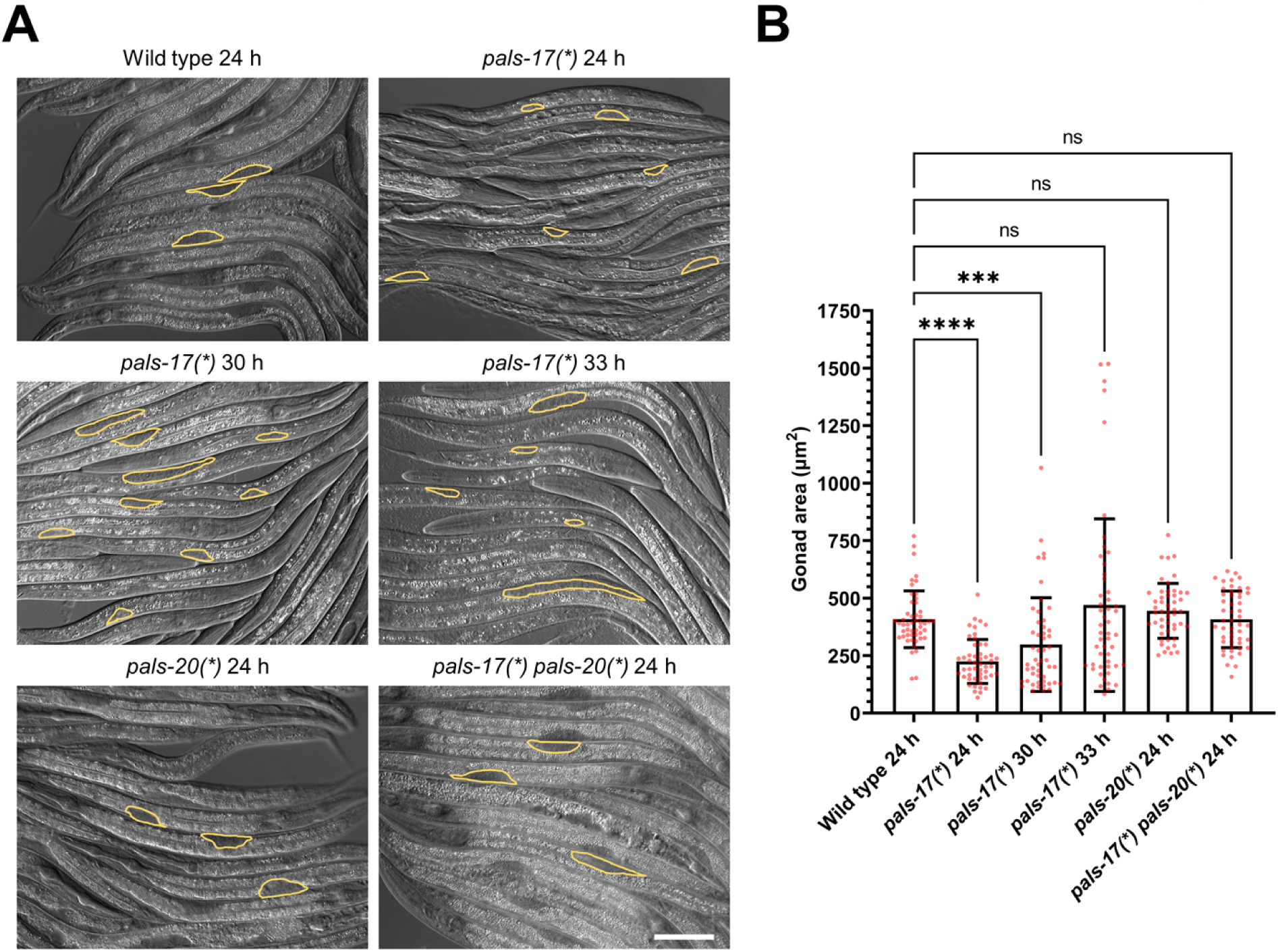
*pals-17* mutants have asynchronous and delayed development. (A) Representative DIC images of *pals-17* and *pals-20* mutants and wild-type control incubated for 24 h at 20 °C, and images of *pals-17* mutants incubated for 30 h and 33 h at 20 °C from L1 stage. The gonads of some animals are outlined with yellow lines. Scale bar, 60 µm. (B) Gonad area measurements. Results shown are the average of two independent experimental replicates, with 25 animals assayed per replicate. Error bars are SD. A Kruskal-Wallis test was used to calculate p-values; *** p < 0.001; **** p < 0.0001; ns indicates no significant difference.

**S4 Fig.**
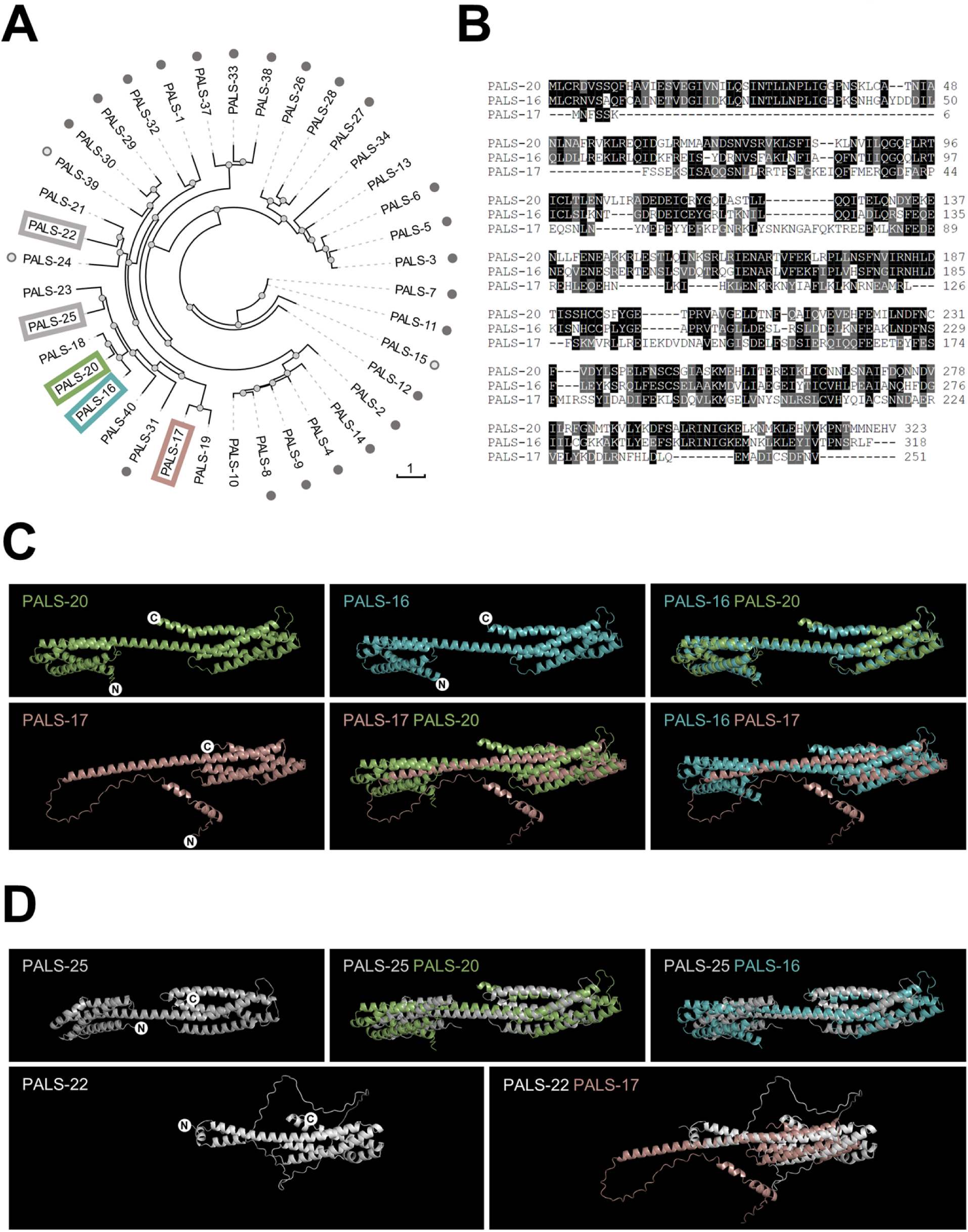
Amino acid sequence analysis and predicted protein structure analysis of PALS proteins. (A) A radial phylogram of the PALS protein family. Dark gray circles indicate PALS proteins whose corresponding mRNA levels are significantly upregulated following IPR activation during microsporidia infection and in *pals-22* mutants (4, 7). Light grey circles label PALS proteins whose corresponding mRNA levels are significantly upregulated only in *pals-22* mutant background. The branch length is indicated by the scale bar. (B) Amino acid sequence alignment between PALS-20, PALS-16 and PALS-17. Black boxes indicate identical amino acids; grey boxes indicate similar residues (defined in Material and Methods). (C, D) predicted PALS protein structures and their overlap. White circles with letters N and C indicate N- and C-terminuses of PALS proteins, respectively.

**S5 Fig.**
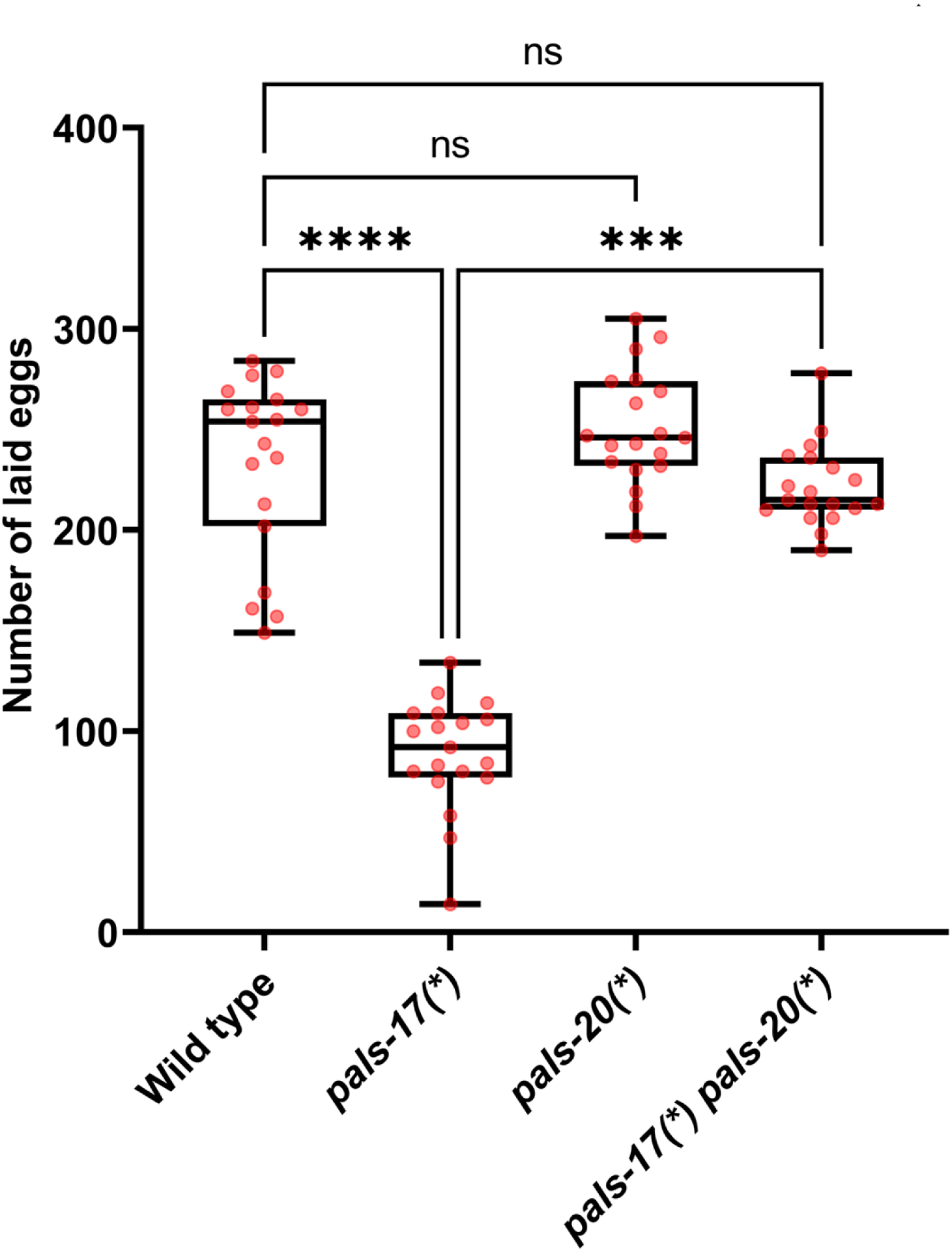
*pals-17* and pals-20 regulate brood size of *C. elegans*. *pals-17* mutants have significantly lower brood sizes, and this phenotype is *pals-20*-dependent. Brood size measurements are shown as a box-and-whisker plot for wild-type, *pals-17(*)*, *pals-20(*)* and *pals-17(*) pals-20(*)* animals. Box lines represent the median values, box bounds indicate 25^th^ and 75^th^ percentiles, and whiskers extend to the minimum and maximum values. Red dots represent individual values for each animal; 19 animals were analyzed for each strain (at least five animals per each of the three experimental replicates). A Kruskal-Wallis test was used to calculate p-values; **** p < 0.0001; *** p < 0.001; ns indicates no significant difference.

**S6 Fig.**
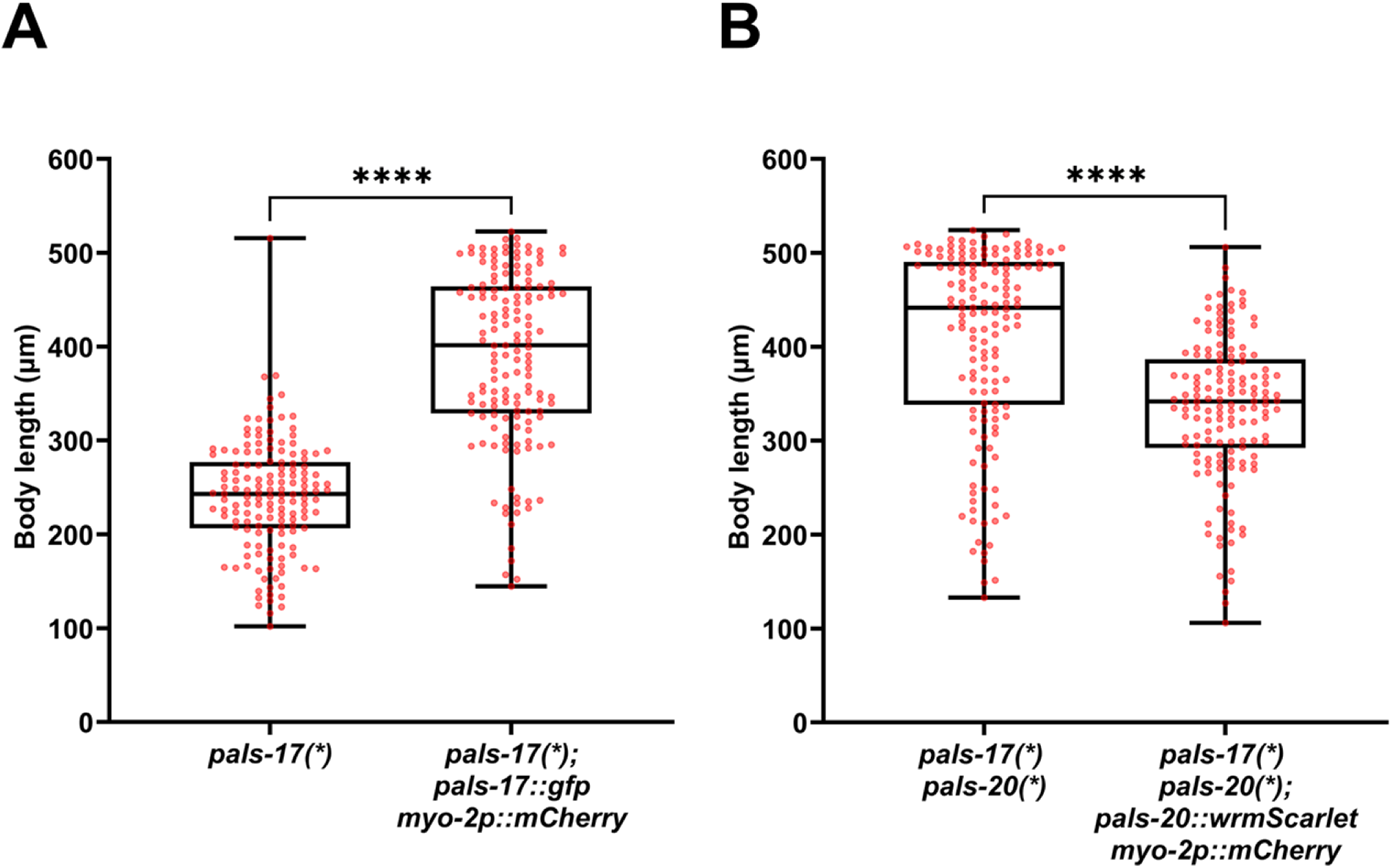
*pals-17* and *pals-20* translational reporters rescue growth phenotypes of *pals-17(*)* and *pals-17(*) pals-20(*)* mutants, respectively. (A, B) Body length measurements are shown as box-and-whisker plots for *pals-17(*)* (A) and *pals-17(*) pals-20(*)* animals (B). Animals expressing *pals-17::gfp myo-2p::mCherry* (A) and *pals-20::wrmScarlet myo-2p::mCherry* arrays (B) as well as their non-transgenic siblings were analyzed. Box lines represent median values, box bounds indicate 25^th^ and 75^th^ percentiles, and whiskers extend to the minimum and maximum values. Red dots represent individual values for each animal; 50 animals per each of the three experimental replicates were analyzed. A Kolmogorov-Smirnov test was used to calculate p-values; **** p < 0.0001.

**S7 Fig.**
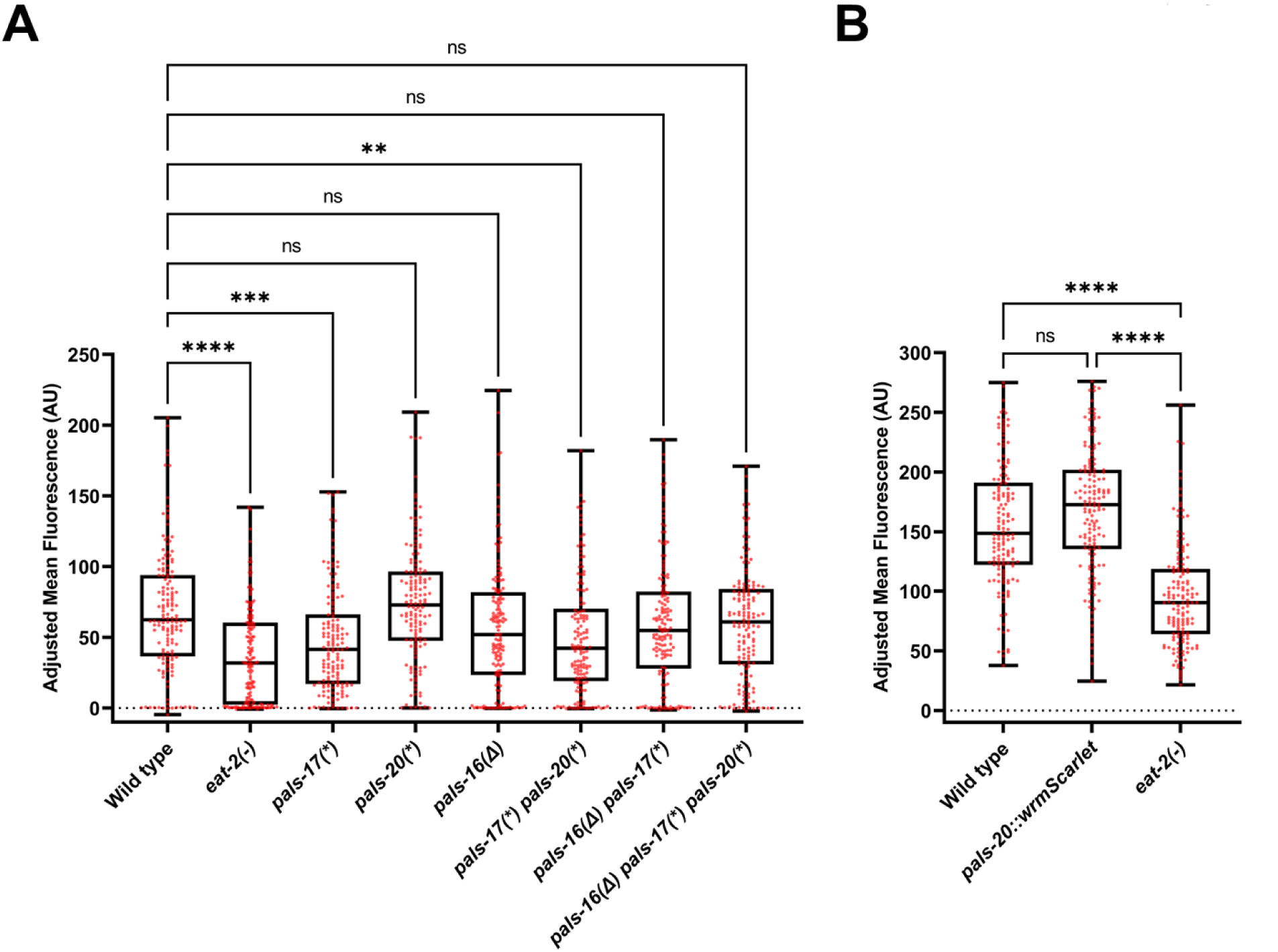
Quantification of fluorescent bead accumulation in *pals-17*, *pals-20*, *pals-16* mutants and control strains. (A, B) Box-and-whisker plots of bead fluorescence levels per animal. Box lines represent median values, box bounds indicate 25^th^ and 75^th^ percentiles, and whiskers extend to the minimum and maximum values. Data from three independent experimental replicates are shown. Red dots represent individual values for each animal; 50 animals were analyzed per strain per replicate. *eat-2(ad465)* mutant was used as a feeding-defective control. A Kruskal-Wallis test was used to calculate p-values; **** p < 0.0001; *** p < 0.001; ** p < 0.01; ns indicates no significant difference. AU = arbitrary units.

**S1 Table. Data shown in graphs in main and supplementary figures**

**S2 Table. Amino acid sequence analysis of PALS-17, PALS-20 and PALS-16**

**S3 Table. Differential expression analysis of RNA-seq data**

**S4 Table. Normalized counts from RNA-seq analyses**

**S5 Table. Full datasets from GSEA analyses**

**S6 Table. List of strains used in this study**

**S7 Table-Primers used in this study**

